# Fitness benefits from co-display favour subdominant male-male partnerships between phenotypes

**DOI:** 10.1101/2022.09.30.510252

**Authors:** James D. M. Tolliver, Krisztina Kupán, David B. Lank, Susanne Schindler, Clemens Küpper

**Affiliations:** Research Group for Behavioural Genetics and Evolutionary Ecology, Max Planck Institute for Ornithology, 82319 Seewiesen, Germany; Department of Biological Sciences, Simon Fraser University, Burnaby V5A 1S6, Canada

**Keywords:** alternative reproductive tactics, Calidris pugnax, lekking, male-male cooperation, reproductive skew, sexual selection

## Abstract

Male-male competition over matings is a key driving force in the evolution of courtship. Typically, competition is an individual affair selecting for dominance and aggression. Yet, some males forgo direct confrontation and improve their reproductive success through cooperation. Occasionally, this leads to specialized alternative reproductive tactics that operate at the intersection of cooperation and conflict. We used a community game model, informed with empirical data derived from previous studies, to examine cooperation dynamics between lekking male ruffs (*Calidris pugnax*) using two different tactics: resident and satellite. Residents defend display courts against other residents on leks. In contrast, satellites forgo court defence and engage in cooperative co-display with selected residents. Co-displaying appears to alter female mate choice, yet the exact mechanism and consequences remain unclear. We modelled individual male mating success as a function of lek size, resident rank, and satellite competitiveness. Our most realistic model assumed that co-display draws copulations from residents proportional to the existing mating skew among them. Under this assumption, all residents benefit from co-display over single display when a satellite is on the lek, except for α-residents co-displaying with the most competitive satellites on large leks. Thus, satellites could nearly always choose their preferred co-display partner, but achieved the highest copulation rates with lower ranking (subdominant) residents on intermediate sized leks. Co-display between the satellite and lower ranking residents reduced the mating skew among residents. However, since copulations for satellites were similar over a range of potential co-display partners, a variety of co-displaying dyads is to be expected, which is consistent with observations in nature. We conclude that, given our model assumptions, co-displaying reduces the impact of male dominance on reproductive success and ultimately alters the course of sexual selection.

## Introduction

Male-male competition over reproduction is typically seen as individual contests over fertilisation opportunities that are won by the most dominant males (Andersson 1994). Under intensive sexual selection, a small number of males then monopolize most of the matings. In such circumstances, however, some males may improve their chances of mating through male-male cooperation and/or developing alternative reproductive tactics.

A prominent form of male-male cooperation is cooperative display (aka co-display) that typically occurs in fishes and birds (Taborsky 1994; Díaz-Muñoz et al. 2014). For co-display, two or more males form temporary partnerships, which may develop into long lasting alliances (Foster 1977; Taborsky 1994; DuVal 2013). These co-displaying males often coordinate their courtship—as well as nest defence in fishes (Taborsky 1994). Co-display may act as an “extended phenotype” increasing one or more of the male(s)’s attractiveness to females (Dawkins 1978). Alternatively, co-display may provide other benefits to females such as protection from harassment during mating (Díaz-Muñoz et al. 2014). Co-displays typically consist of dominant and subordinate males (McDonald and Potts 1994; Krakauer et al. 2005; DuVal 2007). The dominant males mate most often whereas their subordinate partners increase their fitness largely through indirect benefits such as training, subsequent dominance ascension, territory inheritance, or shared genetic relatedness with the dominant male (Díaz-Muñoz et al. 2014). In other cases, the dominant male is unable to maintain complete reproductive control and subordinate males are able to obtain direct benefits, i.e. a share of the matings for themselves (Taborsky et al. 1987; DuVal 2007; Stiver and Alonzo 2013; Hellmann et al. 2015).

Similarly passive and subordinate males can obtain copulations through alternative reproductive tactics (ARTs)—sometimes including co-display (Taborsky 1994; Oliveira et al. 2008). ARTs often occur when sexual selection is strong and territorial males invest heavily in courtship and territory defence leaving themselves open to exploitation by non-territorial males (Jones et al. 2001; Taborsky and Brockmann 2010). Males engaging in ARTs add a novel dimension to the competition for females by engaging in parasitic subterfuge and/or cooperation. Sneaker morphs often parasitize the courtship efforts of territorial males and steal copulations, for example, through female mimicry (Shuster and Wade 1991; Taborsky 1994; Jukema and Piersma 2006). Other ARTs are not fully parasitic and include cooperative elements. Satellite males associate with territorial males and may provide help in the form of territorial co-defence, alloparental care, or co-display in courtship (Taborsky 1994; Widemo 1998a; Stiver and Alonzo 2013; Hellmann et al. 2020). Satellites are rarely able to maintain a territory by themselves, instead they benefit from the courtship efforts and attractiveness of their cooperating hosts. Yet, when it comes to fertilisations, the satellite and its host regularly come into conflict. Since the satellites are already in close proximity, they are able to obtain some fertilisations when females visit, if their dominant partners cannot fully exclude satellites from reproduction (Waltz 1982; Taborsky 1987; Stiver and Alonzo 2013). Satellite males are often formidably prepared for the resulting sperm competition with territorial males, boasting large testicular sizes, highly competitive sperm morphology, and/or unique ejaculate traits (Taborsky 1994; Küpper et al. 2016).

The formation of male partnerships with direct benefits is a strategic process requiring all partners to delicately balance cooperation and conflict over resources (Mesterton-Gibbons et al. 2011 and Oliveira et al. 2008). Theory and empirical evidence suggest that male-male partnerships with direct benefits form when the members can maximize their own reproductive success through the partnership (Noë 1992; Bissonnette et al. 2015). The formation of these partnerships and their composition often depend on the social status of males in the local social environment. For example, high ranking males may be better off without a cooperation partner when competition over resources is low and territories are easily maintained; contrastingly, they may need partners in order to maintain territories or their high social status when competition over resources is intense (Taborsky 1987; Duffy et al. 2007). Similarly, subordinate males may pair up with other lower ranking males to challenge single high ranking males when high ranking males are easily outcompeted (Noë 1992; van Schaik et al. 2004; Bissonnette et al. 2011). Alternatively, subordinate males may seek to associate themselves with high ranking males to benefit from their attractiveness to females and/or to seek protection from other males (Waltz 1982; Oliveira et al. 2002; van Schaik et al. 2004). Such male-male partnerships inevitably alter the mating skew among males (Heinsohn et al. 2000; Taborsky 2008; Bissonnette et al. 2011). The mating skew increases when partnerships include mostly high ranking males and the skew decreases when partnerships include mostly low ranking males. Partnerships between ARTs likely also decrease the mating skew (i.e. when satellites can obtain matings).

The ruff (*Calidris pugnax*) provides an ideal study system to investigate male-male cooperation dynamics between ARTs in variable social environments. Ruff males show an extraordinary behavioural diversity including two displaying morphs, independents and satellites, as well as a parasitic morph called faeders that mimic females. The morphs are genetically determined and exist at relatively stable population proportions (independents ≈ 81%, satellites ≈ 19%, and faeders <1%; Lank et al. 1995; Lank et al. 2013; Küpper et al. 2016; Lamichhaney et al. 2016). Whereas the independent reproductive morph is associated with an ancestral allele, the satellite and faeder morphs are each associated with their own autosomal inversions, which are homozygous lethal (Küpper et al. 2016; Lamichhaney et al. 2016). Independents are aggressive males who compete for resident status on leks (Lank and Smith 1987; Widemo 1998a; Vervoort and Kempenaers 2019). On these leks, residents defend small courts and compete with one another to attract females (Darwin 1871; Hogan-Warburg 1966; van Rhijn 1973; Hill 1991; Widemo and Owens 1995; Widemo 1997).

Residents form a dominance hierarchy where males who are the most aggressive and establish their courts first are the α-males (Widemo 1997). This dominance hierarchy is positively associated with female choice, where α-males attract the highest number of females and receive the highest number of copulations, whereas low-ranking residents are less competitive and attract few, if any, females (Widemo 1997; Widemo 1998b). Lek size also affects the mating competition, as female ruffs prefer to visit and mate on larger leks over smaller ones (Lank and Smith 1992; Höglund et al. 1993). Furthermore, the mating skew among residents decreases with increasing lek size (Widemo and Owens 1995).

Satellites, the other displaying morph, deploy a non-aggressive and non-territorial male ART. They associate with residents, particularly, in the presence of females (Hogan-Warburg 1966; van Rhijn 1991; Widemo 1998a; Vervoort and Kempenaers 2019). Satellite males predominantly visit courts that also receive a large number of visits by females (Hill 1991; van Rhijn 1991; Widemo 1998a). Four potential explanations for this positive association have been proposed (Widemo 1998a). First, satellites may track female movements. Second, satellites might share female preferences for certain residents. Third, females may simply track satellite movements on the lek. Fourth, females might prefer co-display of residents and satellites to single display of residents. The first two hypotheses assume that satellites are reproductive parasites. However, residents not only tolerate satellites, but also actively recruit them onto their courts and achieve higher copulation rates when co-displaying with a satellite (Hogan-Warburg 1966; Shepard 1975; Hill 1991; van Rhijn 1991). Hence, the third and fourth hypotheses, which postulate benefits for residents who share their courts with satellites, are more consistent with empirical observations. In line with this, a previous game-theoretical model predicted that despite the cost of satellite matings, cooperative residents in general will outcompete non-cooperative residents because the co-displaying will attract more female visits to their courts, drawing copulations away from all the other residents (Hugie and Lank 1997). Although this model examined residents who accept/reject satellites for co-display and the optimal frequency of satellites in a population, it did not consider individual variation of payoffs within resident/satellite partnerships.

The resident/satellite partnership is delicate because once a female enters their court, the satellite and resident are in conflict over the opportunity to mate (van Rhijn 1973, 1991). Residents are facultative cooperators that may display alone or with a satellite partner (fig 1A).

**Figure 1.**
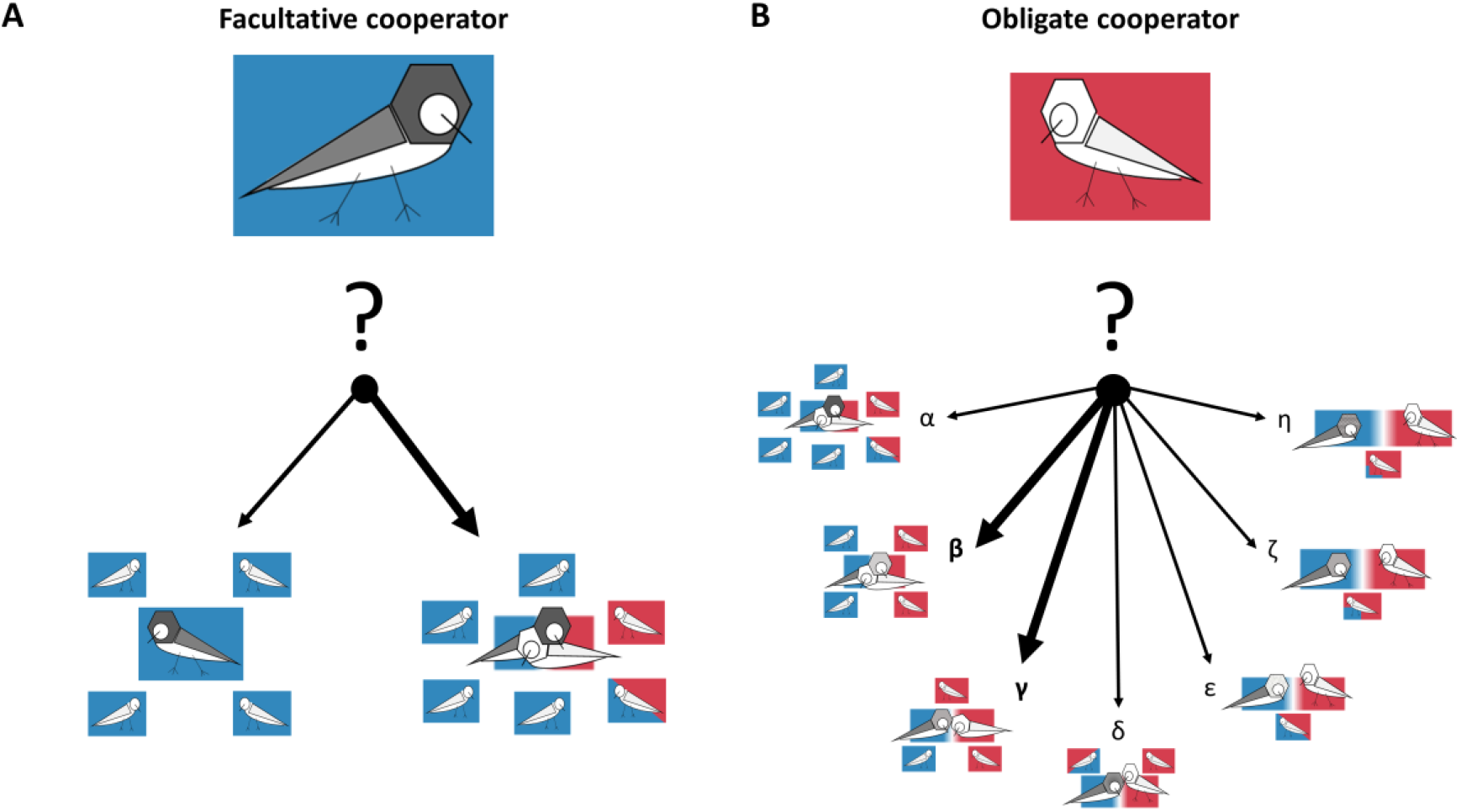
The decision process for male-male co-display differs between facultative cooperators (residents) and obligate cooperators (satellites). Both types of males attempt to maximize their reproductive fitness through cooperation. **A** Lekking residents (dark bird on blue background) should prefer co-display over single display when they can increase their copulation rate (females with blue background) with a satellite (thick arrow). **B** Lekking satellites always require an accepting resident for co-display. Their mating prospects will depend on how many females visit the court and the resource holding potential of their resident partner. If several residents are accepting, satellites should prefer the partner with whom they can maximize their copulation rate (thick arrows and females with red background).

The extent to which particular residents might benefit from co-displaying should influence their willingness to accept a particular satellite partner, but this has not previously been considered. The benefits will depend on social variables including the strength of female preference for co-display, lek size, the male-male rank/competitive abilities of residents and satellites, and finally the individual attractiveness of each partner to a female (Hogan-Warburg 1966; van Rhijn 1983; Hill 1991; Höglund et al. 1993; Widemo and Owens 1995; Widemo 1998a). For their part, satellites are obligate cooperators who should choose the accepting resident partner who maximizes their number of copulations (fig. 1B), which will be a function of social variables such as lek size, their own competitiveness and the competitiveness of their prospective resident partner.

We model a community game exploring cooperation between males through co-display. Community games are derived from economic game theory and particularly suitable to explain the formation of male partnerships, since specific resource holding potentials (RHP) and other individual characteristics can be specified (Mesterton-Gibbons et al. 2011). Our model uses the resident/satellite co-display to examine the consequences of cooperation on male mating skew under a range of naturally observed ruff lek sizes. An important feature of the ruff system is that cooperation only occurs between different male morphs meaning that the actors in our game have specific and fixed strategies. Furthermore, the pronounced hierarchy of the lekking residents and the diversity of lek sizes allows for the examination of costs and benefits of cooperation at the individual level under different social environments. To model female choice, we used literature estimates of resident reproductive skew and copulations per lek (Widemo and Owens 1995). This allowed us to make biologically feasible predictions on the payoffs and rewards of co-display for residents and satellites.

Our model addresses three crucial questions important for understanding male-male cooperation. First, we examine how variation in social environments (lek size and resident rank) influences the propensity for male-male cooperation. We hypothesized that the willingness of residents to co-display with satellites varies with lek size and hierarchical rank. In general, low ranking males should be receptive to co-display, since satellite presence substantially increases the attractiveness of their courts. Similarly, higher ranking residents on smaller leks should be receptive to co-display, since they dominate nearly all the matings and risk losing a substantial proportion of copulations if co-display happens on a rival court. In contrast, high-ranking residents on larger leks may reject co-display, since high competition among residents decreases their ability to monopolize copulations. Second, we examined the satellite’s preference for a specific resident partner. Satellites might maximize their copulation rates by choosing the highest-ranking accepting resident. Alternatively, satellites might prefer low ranking males for a partnership if these low ranking males are less able to monopolize matings within the co-displaying partnership. Here, co-displaying primarily functions as a partnership of lower ranking males against the α-resident on a given lek. Third, we modelled how the most likely co-display partnerships influence the mating skew among lekking residents. Satellite presence should always alter the mating skew between satellites and residents on a lek, but their effect on the skew among residents remains unclear. If co-display is attractive to females, copulations will be re-distributed between resident courts. Finally, we assessed the validity of our model by examining the sensitivity of the predictions to variation in satellite competitiveness and competition between residents.

## Methods

### General model for co-display

We developed an analytical community game to test how male-male co-display arises and is maintained when both males require direct fitness benefits (summarized in Appendix A). Our model consisted of two player classes: dominant and subordinate males. Each player could either perform single display or co-display with the other player type, but not with their own type. Both player classes pursued a strategy where they chose co-display over single display when their fitness payoffs from co-display were equal to or larger than their payoffs when engaged in single display (Mesterton-Gibbons et al. 2011). We assumed that players would always choose partners that would maximize their payoffs when faced with the choice of multiple cooperative partners (“general model of partner choice”, Sachs et al. 2004, Mesterton-Gibbons et al. 2011) because all players were considered to be experienced breeders. This means that we assume our model outcomes represent predictions after either a learning or negotiation process has already occurred and the predictions are average outcomes of these processes.

For co-display between the subordinate male *i* and the dominant male *j*, two factors influenced their respective co-display payoffs *C*. First, we set the synergistic payoff to the co-displaying partnership 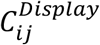. This payoff was the sum of the payoffs of both playersduring single display 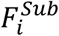 and 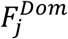, a common assumption in coalition or alliance formation theory (e.g. Mesterton-Gibbons 2011), plus an added benefit for co-display called the “co-display benefit” *F*^*Sub+Dom*^:

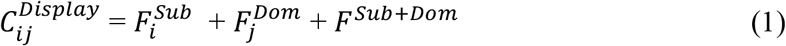

The co-display benefit represented a female preference for co-display over single display. Second, co-display resources were allocated to each co-displaying partner by the dominant male’s RHP 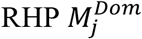:

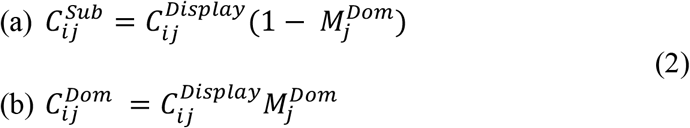

Here 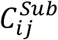, was the fitness payoff to the subordinate male and 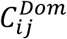 was the fitness payoff to the dominant male from co-display.

We based male co-display between potential partners on a choice function (table A1, numbers 13 – 14). When choosing to co-display with a given male, a male had three options: 1) agree to co-display, 2) ambivalence for co-display, or 3) reject co-display. The choice *A* that each male made was dependent on the potential fitness reward *G* that the male would receive in a co-display. Males would agree to co-display (*A* = 1) when *G* was positive, they would be ambivalent toward co-display (*A* = 0.5) when *G* was equal to zero, and they would reject co-display (*A* = 0) when *G* was negative. Co-display occurred when both males agreed to co-display, when one male was ambivalent and the other agreed to co-display, or when both males were ambivalent. The fitness reward 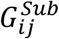 of the subordinate male was the difference between his co-display payoff 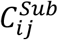 and his single display payoff 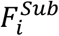 (equation 3a). Similarly, the fitness reward 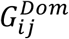 of the dominant male was the difference between his co-display payoff 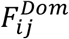 and his single display payoff 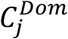 (equation 3b).

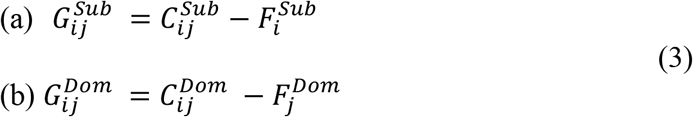

Thus, the decisions of the subordinate and dominant males to co-display can be determined when both the total fitness payoff available to a co-display 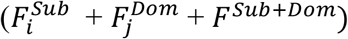 and the RHP of the dominant male 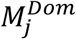 are known.

### Modelling resident/satellite co-display in ruffs

We applied our model, along with its assumptions to lekking ruffs to predict when cooperation between satellites and residents should arise (fig. 1; table 1; Appendix B in *Ruff model assumptions*). Ruffs fit well into our assumption that players are experienced breeders (i.e. the full information and explicit RHP framework) because lekking males are usually older and experienced males that have high lek fidelity across years (van Rhijn 1991). When ruff males co-display, residents are the dominant and satellites the subordinate partner (Hogan-Warburg 1966; van Rhijn 1973). Within the model, we limited the number of satellites per lek to one to restrict model complexity. The restriction still covers the dynamics on most naturally occurring leks as the average number of competitive satellites per lek is about one for most lek sizes, and non-competitive satellites generally do not obtain copulations (van Rhijn 1991; Höglund et al. 1993; Vervoort and Kempenaers 2019). In addition, satellites are rarer and spend less time on leks when compared to residents (Widemo 1998a; Vervoort and Kempenaers 2019).

**Table 1.**
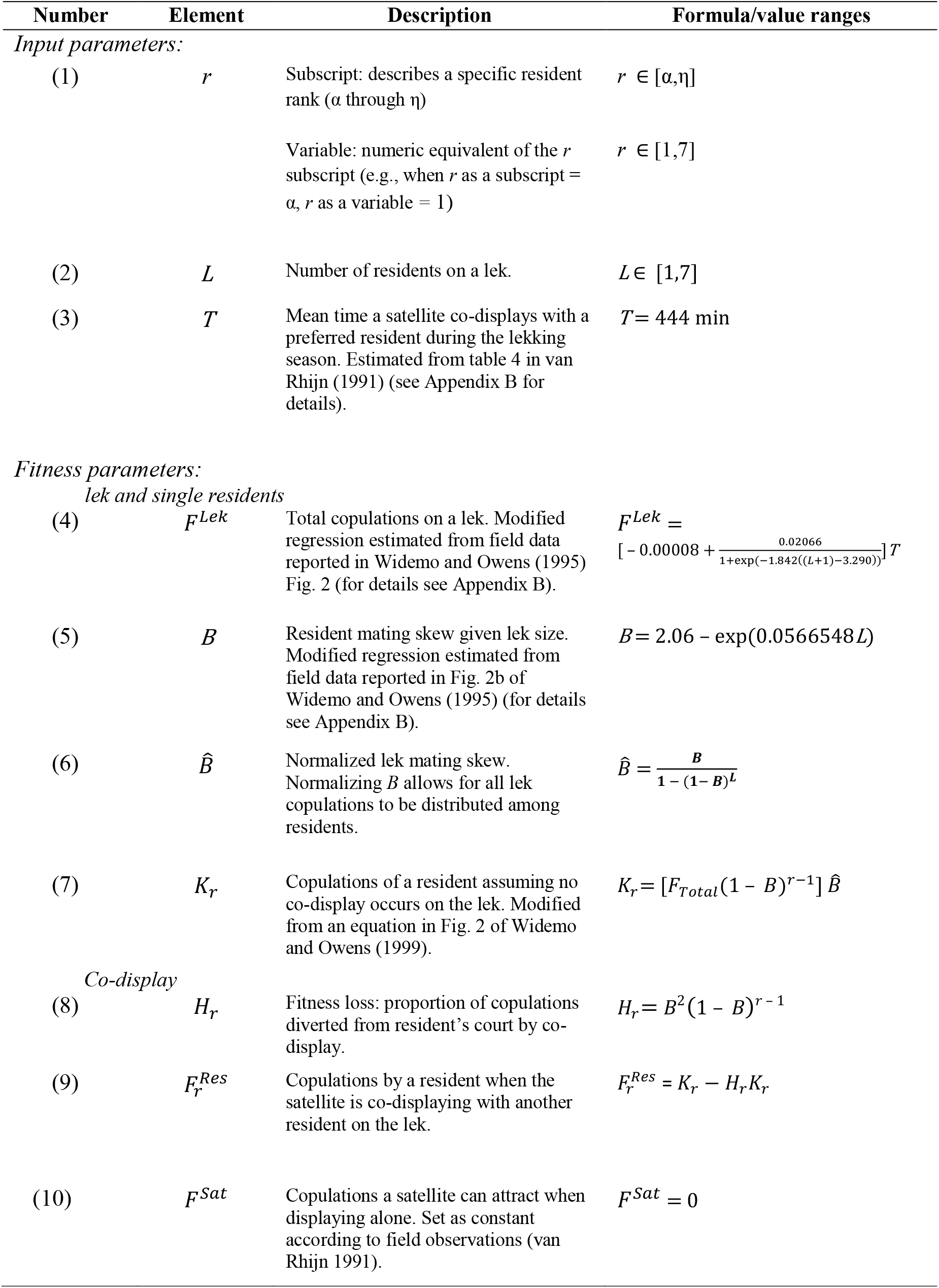
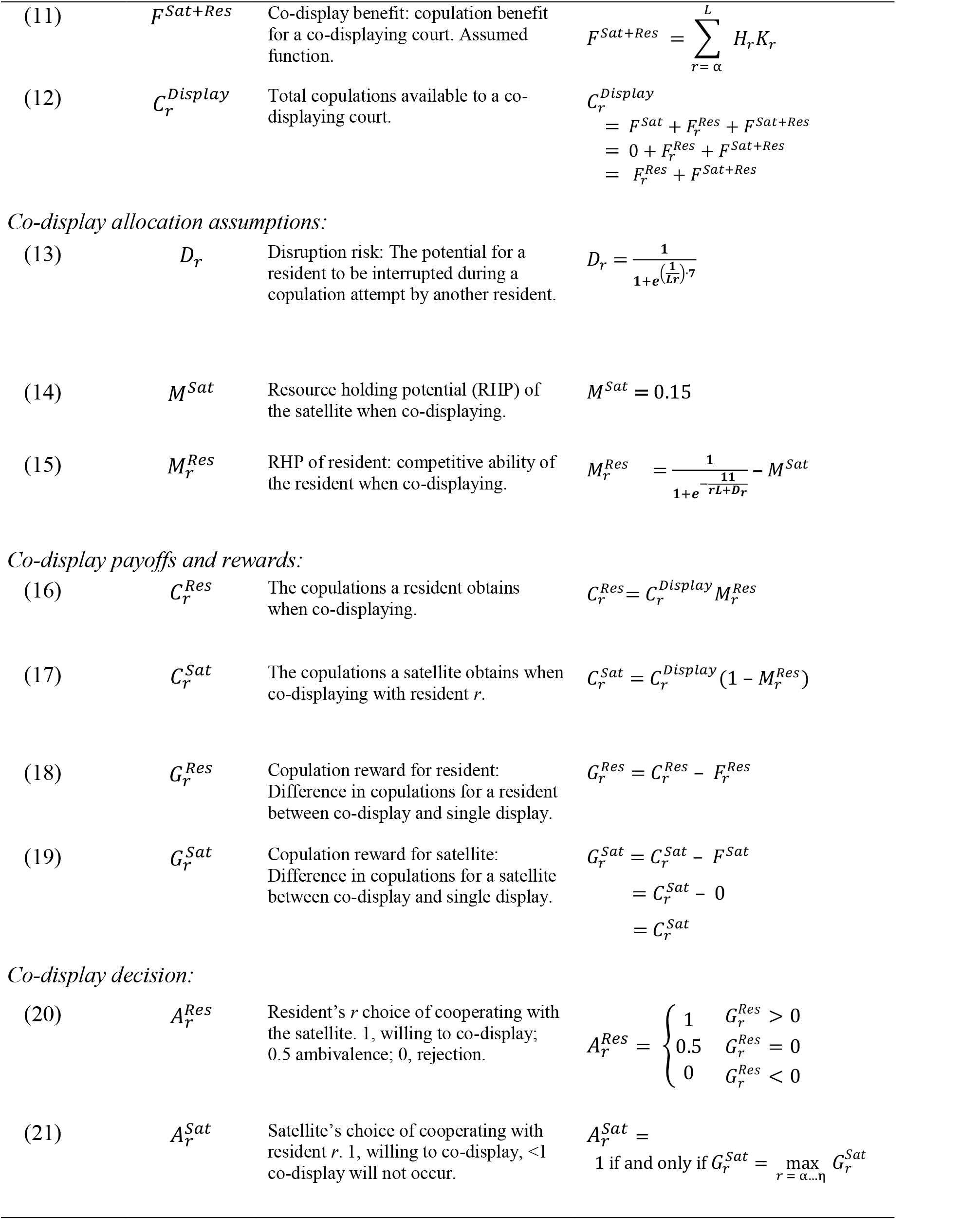
Notation and definitions of ruff co-display model. All functions in table are assumed unless stated otherwise in the description.

We adjusted the parameters of the general model described above to fit ruffs. For individual residents, the index *r* described the hierarchical rank in descending dominance from α = 1 to η = 7 (table 1, number 1). 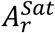 referred to the satellite choice of resident *r* for co-display. 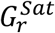 specified the potential satellite copulation reward when co-displaying with resident *r*. 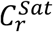 specified the realized satellite copulation payoff when co-displaying with resident *r*.*F^Sat^* indicated the payoff of a satellite during single display; with the analogous parameter 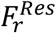 for a resident.

Satellites had a more specific strategy than the subdominant male in the general model because they rarely obtain copulations outside of a co-display (van Rhijn 1991) and consequently we always set *F*^*Sat*^ to zero. Hence, the satellite always chose the accepting resident with whom he could maximize his payoff during co-display, given by:

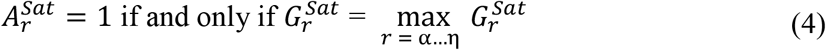

Residents had a similar strategy as dominant males in the general model; they co-display with a satellite whenever 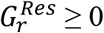 (table 1, number 20).

### Parameterization of the ruff co-display model

From published data, we obtained biologically meaningful estimates of female choice for two parameters: copulations payoffs for a single resident 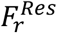 and the resident/satellite co-display benefit *F*^*Sat+Res*^. We parameterized resident copulation success on leks using observed copulations rates from Widemo and Owens (1995), taking into account hierarchical rank and lek size as these two social environmental factors correlate with resident mating success (Höglund et al. 1993; Widemo and Owens 1995; Widemo 1998b). We estimated these copulation rates with two regressions from Widemo and Owens (1995). The response variables for these regressions were total number of copulations per lek and the resident mating skew, whereas the predictor variable for both regressions was the number of residents per lek (table 1; fig. B1; Appendix B in *Model parameterization*). Widemo and Owens (1995) only used copulation rates of residents to estimate these regressions; whereas information on satellite presence and their respective influence on female choice (i.e. copulation rates) on different leks were not provided.

To overcome this lack of data, we assumed that the presence of a satellite on a lek was equivalent to having another resident on that lek when estimating total copulations per lek. That is, we simply added one to the predictor variable, residents per lek, *L* (table 1, number 4). This assumption is supported by field observations that, when time on a lek is controlled for, mating success per male does not differ as a function of morph (Vervoort and Kempenaers 2019). We multiplied the total number of copulations by the mean time (in minutes) that a satellite spends with a preferred resident during the peak lekking season (Appendix B in *Model parameterization*). We then allocated those copulations to each resident using a decay function (table 1, numbers 4 – 7). In the absence of co-display, the decay function followed the resident hierarchy and the rate of decay was equal to the normalized resident mating skew from Widemo and Owens (1995). This represents the female choice in resident (i.e. copulations per resident) as copulations per resident is correlated with dominance rank in ruffs (Widemo 1997). We assume these copulations, *K*_*r*_, represent the payoff each resident would receive if a satellite were on the lek but engaged only in single display and not in co-display.

As noted above satellites would never engage in single display (*F*^*Sat*^ = 0). Thus, we modified *K*_*r*_ to account for the observed positive correlation between satellite presence and female visitation rates on a resident’s court. We assumed that co-display would redistribute a fraction of copulations from all courts of a lek to the court with co-display (Appendix B in *Fitness loss assessment*; table B1; figs. B2 – B3). This increase in copulations on a co-display court was the co-display benefit *F*^*Sat+Res*^, given by:

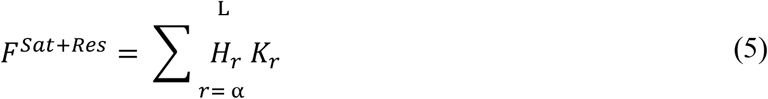

The fraction of copulations taken from each resident’s court was the fitness loss *H*_*r*_. Thus, 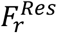 the payoff for a resident during single display was the difference between *K*_*r*_ and the amount of fitness lost to a co-display if the resident rejected the satellite (equation 6).

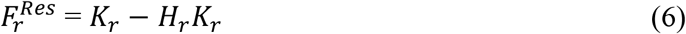

This assumption means that when co-display occurred on a lek, copulations were taken from each resident and moved to the co-display court via the equations 5 and 6. Further, it implies that all residents on a lek with co-display attract fewer copulations than they would if all the males on the lek were only residents. This is because we assumed that the total copulations on a lek only depends on the number of males on a lek, regardless of morph. Therefore, a resident decides to reject or accept a satellite for co-display by balancing his fitness loss, *H*_*r*_, if rejecting the satellite against his RHP 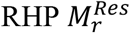 if accepting the satellite (Appendix C, *Limits of Resident co-display choice*).

As the exact mechanism and magnitude of *H*_*r*_ is unknown, we assumed that most copulations would be drawn from high-ranked residents and the fewest would be drawn from the low-ranked residents (“Skew” fig. B2*C*; table B1; table 1 number 8). We examined three other hypothetical scenarios: “Null”, “Uniform Proportion”, and “Reverse Skew” (figs. B2 – B7; table B1, numbers 3, 4, and 6) to see which scenario produced biologically realistic outcomes. Our model predictions were highly sensitive to these alternative scenarios. However, only the “Skew” scenario predicted patterns of co-display with satellite copulation rates comparable to those observed in nature (ca. 10%; fig. B7; Höglund et al. 1993; Widemo 1998a) for all lek sizes. Hence, we present the “Skew” scenario in the main text, as it was the most biologically realistic model, (fig. 2; figs B4*C* – B7*C*), but provide the corresponding annotations and results of the other scenarios in the Supplement (figs. B4 – B7).

**Figure 2.**
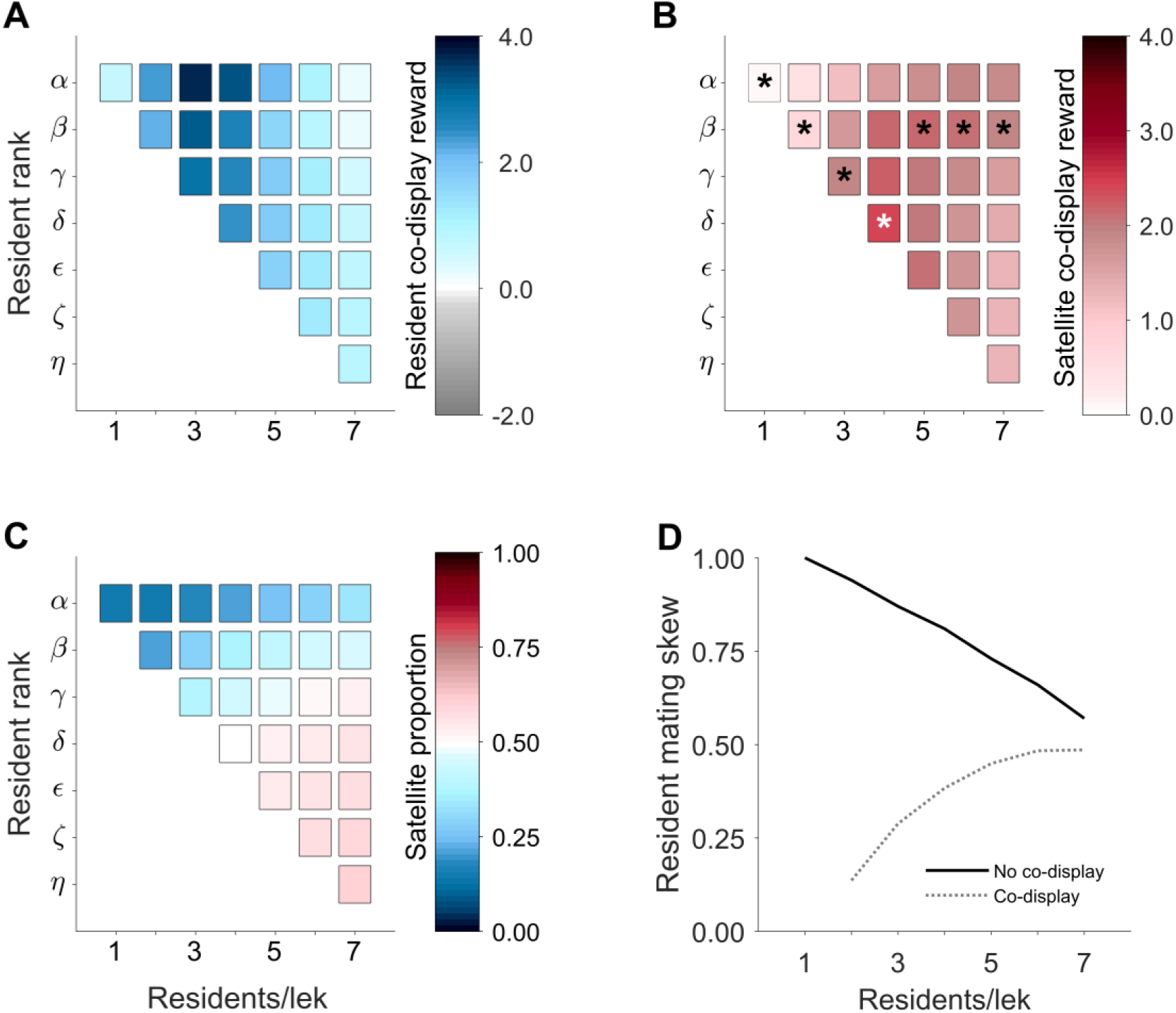
Predicted fitness consequences during co-display between residents and satellites according to lek size (residents per lek) and hierarchical rank of residents under the “Skew” scenario. **A** Copulation rewards from co-display for residents. All residents were willing to co-display with the satellites because resident rewards are always greater than zero. **B** Copulation rewards for the satellite from co-display according to the rank of its resident partner. Asterisks indicate preferred satellite choice at a given lek size. Overall, satellites achieve the highest copulation reward when paired with δ-residents at *L* = 4 (white asterisk). **C** Proportion of copulations for each partner of a co-display unit. Satellites gain the highest proportion of copulations when co-displaying with low ranking residents on large leks. **D** Mating skew among residents with or without co-display. Shown is the co-display with the preferred partner of the satellite at a given lek size taken from **B**; these partnerships always reduces the mating skew among residents at a given lek size.

Copulations on courts with co-display were allocated between the satellite and the resident according to 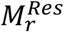, which represents both the competitive abilities of individual residents and satellites, and the female choice towards either the resident or the satellite (fig. B8*A*). 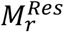 was assumed to be influenced by the threat of copulation interruptions by other residents. Residents frequently interrupt copulation attempts of their neighbours; these interruptions are especially common for residents on large leks and/or of low rank (van Rhijn 1991; Widemo 1997; Widemo 1998b). Consequently, we defined 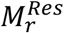 as a function of the lek size *L*; the rank of the resident *r*; the “disruption risk” *D*_*r*_, which refers to the threat of a copulation interruption. We also assumed that the RHP of the satellite *M^Sat^* (i.e. the attractiveness of the satellite to females and the satellite’s ability to interrupt their resident partner’s copulations) influenced 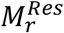 (equation 7).

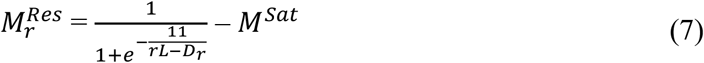

Here we assume that 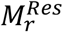 has a range from 0.35 (a η-resident on the largest lek size; *L* = 7) to 0.85 (an α-resident on the smallest lek size; *L* = 1) when paired with a satellite of intermediate competitive ability (*M*^*Sat*^ = 0.15; fig. B8*A*). We assume that *D*_*r*_ increases with lek size but decreases with resident rank and ranges from *D*_*r*_ = 0, on the smallest lek size, to *D*_*r*_ = 0.45 for a η-resident on the largest lek size (table 1, number 13; fig. B8*B*). This form of mechanistic bargaining emphasizes that the resident’s social environment drives the resource allocation rather than the actions of rational actors, unlike other possible allocation approaches (i.e. transferable utility models; Mesterton-Gibbons et al. 2011), and fits into our assumption that negotiation and/or learning has already taken place.

We used the model to predict the pattern of male-male cooperation across different hierarchies and lek sizes, the preferred satellite choice based on their rewards, and the consequences of co-display on the mating skew among residents.

### Sensitivity analysis

Because of a lack of empirical data for 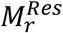 and *D*_*r*_, two fundamental parameters of our model, we examined the sensitivity of our model predictions to perturbations of these parameters (Appendix D). We varied the range of 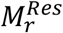 assuming two additional classes of satellite: weak and strong competitors (table D1 and fig. D1*A – C*). We assumed weak satellites would have a RHP equal to zero, *M*^*Sat*^ = 0, with the resulting resident RHP ranging between 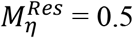, on the largest lek size, *L* = 7, to 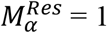 on the smallest lek size, *L* = 1.In contrast, strong satellites could obtain at least 30% of all copulations within a co-display, *M*^*Sat*^ = 0.3, with resident RHP ranging between 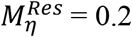, on the largest lek size, and 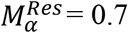, on the smallest lek size. Similarly, we varied the ranges of *D*_*r*_, to model different levels of competition on leks (fig. D1*D – F*). In addition to the intermediate competition scenario, we modelled high competition (*D*_*α*_ = 0.5 on the smallest lek to *D*_*η*_ = 0.95 on the largest lek) and low competition (*D*_*α*_ = 0 on the smallest lek to *D*_*η*_ = 0.25 on the largest lek). Finally, we examined the average change in model parameters given a 0.1 increase in 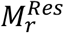 using linear regressions. For these regressions, we used model predictions and coefficients calculated from all combinations of lek size (1 – 7), rank (α – η), satellite strength (weak, medium, strong), and competition levels (low, medium, high). We used MatLab R2017A for all our model predictions (https://github.com/ruffresearcher5/co-display_in_lekking_ruffs).

## Results

### Co-display

In our model, we first investigated which residents agreed to co-display with satellites on leks. In the “Skew” model with a medium strength satellite, male-male cooperation was universal; all residents, regardless of rank and lek size, were willing to co-display with satellites (fig. 2*A*). The other scenarios predicted that resident acceptance of satellites for co-display depended on the combination of rank and lek size (fig. B4 – B5). The copulation reward varied with social environment, i.e. rank and lek size (fig. 2*A* – *B*; B4 – B5). In the “Skew” model, the expected copulation reward from co-display for both residents and satellites was highest at intermediate sized leks (fig. 2*A – B*). However, whereas among residents, α-residents on intermediate sized leks were predicted to receive the highest rewards from co-display, the overall preferred satellite choice for co-display was the lowest ranking resident at a lek size of four (fig. 2*B*). The importance of the resident rank on the satellite reward decreased with increasing lek size, as shown by the reduction in absolute differences in co-display reward. The predicted satellite preference of residents was not the same as the resident with whom the satellite gained the highest proportion of copulations. Although satellites could always claim the highest proportion of copulations when co-displaying with low ranking males (fig. 2*C*), their preferred choice depended on lek size. On large leks (*L* = 6 – 7), satellites received the highest reward when co-displaying with high ranking residents, whereas on intermediate sized to small leks (*L* = 2 – 4) co-displaying with low ranking males provided the largest copulation reward for the satellite (fig. 2*B*).

Co-display always reduced the mating skew among residents in the “Skew” model (fig. 2*D*, B*6*). The reduction was most pronounced at smaller leks, and the impact of co-display on the mating skew became weaker with increasing lek size.

### Model sensitivity

Our model predictions were sensitive to changes in 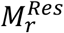 (fig. D2) but not *D* (fig. D3). Increasing satellite competitiveness reduced 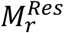, resulting in high ranking residents on large leks becoming more likely to reject satellites. Satellite competitiveness did not influence the satellite’s preferred lek size, but changed their preferred resident partners on larger leks with stronger satellites preferring higher ranking residents (fig. D2*B*). Changes in satellite RHP did not influence the general impact of co-display on resident mating skew. Variation in *D*_*r*_ did not alter the acceptance rates of residents nor the mating skew among residents and only altered the satellite’s preferred resident partner on a lek size of 7 (fig. D3). Although we acknowledge that the influence of *D*_*r*_ on 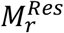 was limited (fig. D4). Examining the consequences of changing the RHP of the co-displaying partners, we found that on average, a 0.10 increase in 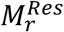 increased the resident copulation rewards from co-display by 0.34 (fig. D5*A* – *B*). Similarly, the proportion of copulations received by the resident within a co-display had a 1:1 increase with 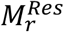 (fig. D5*C*). The resident mating skew resulting from co-display was not affected by changes in 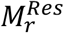 (fig. D5*D*).

## Discussion

Cooperation is considered to be a highly successful strategy for less competitive males to gain reproductive success in an extremely competitive mating environment (van Schaik et al. 2006; Bissonnette et al. 2011). However, these male-male partnerships entail a delicate balance between cooperation and conflict, especially when all cooperating males pursue direct benefits. Therefore, it is essential for males to choose carefully their cooperative partner within variable social environments. Yet, surprisingly, the influence of social characteristics, such as behavioural strategy and hierarchical rank of the potential cooperative partner and/or group size, remain understudied at the individual level (Enigk et al. 2020, Hellmann et al. 2020, Kawazoe 2020; reviewed by Taborsky 2009 and Hager and Jones 2009). Here we used a community game model to investigate cooperative partner choice between co-displaying territorial residents and non-territorial satellites in the lekking ruff.

An earlier game theoretical model for the evolution and maintenance of the resident/satellite system predicted that co-display between satellites and residents would outcompete single display by residents and become a fixed feature in ruff populations (Hugie and Lank 1997). This explained why co-display between residents and satellites is ubiquitous throughout ruff populations and helps account for the maintenance of the satellite allele despite an intrinsic genetic handicap (Küpper et al. 2016). Building on the earlier model, we investigated the variation in fitness benefits of co-display for both residents and satellites with respect to the lek size, social status and competitiveness of potential partners. Our addition of individual RHPs added a conditional tactical component to the universal preference of residents for co-display predicted by Hugie and Lank (1997). That is, our introduction of variation in RHP resulted in the net loss of payoffs for α-residents on the largest leks who co-displayed with highly competitive satellites (D2*A* – *B*). These α-residents should reject satellites whereas all other residents, across different lek sizes and ranks, who should accept satellites, regardless of their RHP, because their payoffs from co-display exceeded the payoffs of single display in presence of a satellite.

Despite our model predicting net positive fitness rewards from most co-displays, the rewards for satellites and residents differed substantially with respect to lek size and resident rank. Overall, our model predicted the highest gains in copulations for cooperative residents and satellites at intermediate sized leks. Consistent with this, field observations show that satellites are most successful on intermediate sized leks (*L ≈* 3.5, Höglund et al. 1993).

Our model predicted that, in most cases, the satellite gained the highest benefit from co-displaying with a sub-α resident. Because of their scarcity and the almost universal willingness of residents to co-display, satellites could choose the resident partner with whom they maximized their reproductive success. For *L* > 1, α-residents were never the preferred choice for co-display because high ranking residents monopolize most of the copulations on a co-displaying court. On larger leks (*L =* 5 – 7), the predicted preference of the satellite switched from the lowest ranking resident to the ß-resident. On small to intermediate leks, our model predicted that satellites preferred to co-display with the lowest ranking males.

However, the cooperation reward predicted on these leks was highest for α-residents, resulting in a stronger conflict of interest among residents and between the satellite and the α-resident, which may affect the formation of co-displays. It is conceivable that α-residents may attempt to bully satellites into co-displaying with them. Assuming our parameterization of RHP reflect those in wild ruffs, such bullying and attacking co-displays should be most effective on small leks where the α-resident has the highest RHP and differences in competitive ability among residents are most pronounced. This would mean that low ranking residents might not be able to sustain co-display on their court (except for satellite choice on the lek size of 7). Note that we varied disruption risk in our sensitivity analysis and found little to no influence on co-display formation. However, this accounted only for competitive interactions between residents, but not the hypothesized antagonistic interactions between α-residents and satellites.

Our predictions about general acceptance of co-display by residents, when satellite RHP was weak to intermediate, is consistent with observations of labile and short lasting co-displaying, with satellites frequently changing resident partners on or between leks (Hogan-Warburg 1966). This lability and the mobility of satellites may also help them to evade bullying attempts by dominant α-residents. Occasionally, residents have been observed to reject co-display (Hogan-Warburg 1966; van Rhijn 1991; Widemo 1998b), which, according to our model should preferably happen by high ranking residents on large leks when facing strong satellites. In other systems with male-male cooperation, partner choice of subordinate males is often limited to other low or similarly ranked males (Bissonnette et al. 2011; Young et al. 2014). Even in species with ARTs, subordinate male satellites can be forced out of partnerships by the cooperating dominant male (Hellmann et al. 2020).

Natural ruff leks are highly dynamic and variable in their composition (Lank and Smith 1987). As empirical studies have yet to address many of these complexities, our model operated with a number of simplifying assumptions. First, we considered only a single satellite per lek. Multiple satellites can attend leks at the same time (van Rhijn 1991; Höglund et al. 1993; Vervoort and Kempenaers 2019). However, a recent study of five ruff leks in Norway found that “central” satellites, i.e. those satellites that engage in co-display and obtain the vast majority of satellite copulations, occurred at a mean number of one per lek (Vervoort and Kempenaers 2019), meaning that our model would be representative of effective mating satellite presence on these natural ruff leks. Second, the satellite and residents in our model were aware of each other’s RHP. This assumption is supported by the fact that resident males are typically older and experienced males that return to the lek site and will have built up knowledge on their direct competitors. In younger males, building up this knowledge might involve a learning process and/or iterative negotiation. It is currently not known whether and to what extent satellites or residents negotiate over copulations, but male ruffs have extraordinary individual variation in their elaborate breeding plumages that would facilitate individual identification and reciprocity (Dale et al. 2001; Lank and Dale 2001). Future models may simulate a learning process or an iterative negotiation between males using our model’s predicted rewards as starting values. Alternatively, controlled experiments involving naïve residents and satellite combinations may shed light on how satellites and residents learn their preferred co-display partners.

As the satellite effect on female choice is not fully understood (Widemo 1998a), we considered multiple scenarios for how satellite presence might affect the distribution of matings on a lek (Appendix B in *Model parameterization*, subsection *Fitness loss assessment*). Overall, the “Skew” scenario, where female choice for co-display is leveling the mating skew when satellites are paired to a lower ranking resident, is most consistent with field data (Fig. B7; Appendix B in *Model parameterization*, subsection *Fitness loss assessment*). Under this scenario, satellites achieved realistic copulation rates (≥ 10%) and co-display was viable for all lek sizes and across all resident ranks (Höglund et al. 1993; Widemo 1998a). Alternative parametrizations where satellites either pursued a purely parasitic strategy (“Null” scenario) or where co-display with high ranking males increased the mating skew among residents (“Reverse skew” scenario) made co-displaying very unlikely (fig. B4). Similarly, assuming an equal proportional loss of copulations for all single residents (“Uniform” scenario), meant that co-displaying would most likely be observed on small to intermediate sized leks or with low ranking males (fig. B4*B*), as satellites would only then obtain their necessary share of copulations (fig. B7*B*; D6). Therefore, even though these different scenarios demonstrate our model predictions are sensitive to the parameterization of the fitness loss, our choice in parameterization is the most consistent with available data.

Female preference for co-displaying could limit lek size. Ruff leks tend to be smaller than those of most other lekking birds (Höglund and Alatalo 1995). Although females of many lekking species, including ruffs, often prefer to mate on larger leks (Alatalo et al. 1992; Lank and Smith 1992; Höglund et al. 1993; Kokko 1997), ruff leks rarely have more than seven residents (Lank and Smith 1987; Höglund et al. 1993; Widemo and Owens 1995; Vervoort and Kempenaers 2019). One explanation for this is that the marginal benefits for prospecting residents decrease with lek size (Widemo and Owens 1995, 1999). Our model also predicts a decrease of benefits for the subordinate partner, here the satellite, as lek size increases. In the wild, satellites have the highest per capita reproductive success on intermediate sized leks (*L* ≈ 3.5; Höglund et al. 1993). However, this could also be a consequence of larger leks having multiple satellites who then compete over co-displaying partners. This should lead to reduced benefits for each co-displaying unit, which warrants further investigation.

Understanding the ultimate consequences and constraints of partner choice provides key insights into the evolution and maintenance of male-male cooperation (Noë 1992; McDonald and Potts 1994; Hellmann et al. 2020). Our model predicts that satellites will prefer subdominant residents over α-residents for co-display. This predicted choice by the satellite can alter the mating skew on leks profoundly. Two mechanisms are responsible for this. First, in the model, the satellite itself obtained a share of the matings reducing the mating skew among all males. Second, since in the model co-display typically occurred between satellites and lower ranking resident males, the resulting skew among the residents also decreased. Previously, it has been shown that sneaking ARTs reduce the variance of male mating success in fish with male care (Jones et al. 2001). Here we demonstrate that a cooperative ART can also reduce the mating skew in a system with strong sexual selection.However, whereas sneaking ARTs usually steal copulations, satellite ruffs divert copulations to the court of their cooperating partner. Thus, female preference for co-display may alter the trajectory of male-male competition and, therefore, sexual selection profoundly, as more cooperative and less aggressive males will become more successful.

## Conclusions

We developed a community game to study cooperation and partner choice with direct benefits in a highly competitive environment. The model predicts that most territorial males prefer co-display over single display. Satellite males are expected to avoid co-display with α-males and prefer instead sub-α males in intermediate competitive environments to maximize their own reproductive success. Co-displaying reduces the mating skew not only through benefitting satellites, but also by reducing the variance in mating success between territorial males. Hence, female preference for co-display profoundly alters the course of sexual selection and affects selection on male behavioural traits such as aggression and cooperative ability.

## Acknowledgements

We thank R. Vervoort for discussing the dynamics of co-display in ruffs. J.D.M.T. K.K. and C.K. were funded by the Max Planck Society, and D.B.L. by the National Science and Engineering Research Council of Canada. We thank E. Akçay, D. Dechmann, L. J. Eberhart- Hertel, W. Forstmeier, L. M. Giraldo Deck, K. Safi, M. Küblbeck, J. L. Loveland, C. Riehl, and M. Taborsky for their comments on earlier drafts of this manuscript.

