## Appendices for "Fitness benefits from co-display favour subdominant male-male partnerships between phenotypes"

### Appendix A

#### Assumptions of general male-male co-display model

1. Total fitness must be $\geq 0$
2. Players are unrelated and cooperate for co-display in pairs
3. Cooperative partners are experienced males, they know the absolute resource holding potential of other players. Results of the model are therefore assumed to be the stable average outcomes of a learning/negotiation process.
4. Cooperation is categorical and not continuous (i.e. individuals will either be rejected, be received with ambivalence, or accept by potential co-display partners).
5. Both cooperators must either accept or be ambivalent to a potential cooperator for co-displays to form
6. The synergistic resources available to the co-displaying partnership consists of the fitness obtainable by each co-display partner alone and the fitness that is only available to the co-display partners through some co-display benefit
7. Each partner has their own competitive ability (resource holding potential) which is influenced by the individuals quality (influenced by social environment and genetics) and the competitive ability of the cooperative partner
8. A dominant and subordinate male will only co-display if each male's reward from co-display is equal to or greater than their single display reward or co-display with a different male
9. Cooperators maintain the cooperation in spite of conflict during a constant time period

Preprint

Table A1. Notation and definitions of general male-male co-display model.

| Number | Element | Description | Formula/value ranges |
|---|---|---|---|
| *Indices:* | | | |
| (1) | $i$ | Index of the subordinate male. | unique ID |
| (2) | $j$ | Index of the dominant male. | unique ID |
| (3) | $ij$ | Index of a co-display between subordinate male *i* and dominant male *j*. | unique ID |
| *Fitness assumptions:* | | | |
| *Single display* | | | |
| (4) | $F_i^{Sub}$ | Fitness obtainable by subordinate male *i* when displaying alone. | $F_i^{Sub} \in$ [0, max. fitness of *i*] |
| (5) | $F_j^{Dom}$ | Fitness obtainable by dominant male *j* alone. | $F_j^{Dom} \in$ [0, max. fitness of *j*] |
| *Co-display* | | | |
| (6) | $F^{Sub+Dom}$ | Fitness benefit for a co-display between the subordinate and dominant male. | function of available fitness |
| (7) | $C_{ij}^{Display}$ | Synergistic resources available to a co-display unit of males *i* and *j*. | $C_{ij}^{Display} = F_i^{Sub} + F_j^{Dom} + F^{Sub+Dom}$ |
| *Co-display allocation assumption:* | | | |
| (8) | $M_j^{Dom}$ | Resource holding potential of the dominant male *j*. | $M_j^{Dom} \in [0,1]$ |
| *Co-display payoffs and reward:* | | | |
| (9) | $C_{ij}^{Sub}$ | Payoff for subordinate male *i* from co-display with dominant male *j*. | $C_{ij}^{Sub} = C_{ij}^{Display}\,(1 - M_j^{Dom})$ |
| (10) | $C_{ij}^{Dom}$ | Payoff for dominant male *j* from co-display with subordinate male *i*. | $C_{ij}^{Dom} = C_{ij}^{Display} M_j^{Dom}$ |
| (11) | $G_{ij}^{Sub}$ | Fitness reward for subordinate male *i* from co-display with dominant male *j*. | $G_{ij}^{Sub} = C_{ij}^{Sub} - F_i^{Sub}$ |
| (12) | $G_{ij}^{Dom}$ | Fitness reward for dominant male *j* from co-display with subordinate male *i*. | $G_{ij}^{Dom} = C_{ij}^{Dom} - F_j^{Dom}$ |
| *Co-display decision:* | | | |
| (13) | $A_{ij}^{Sub}$ | Decision by subordinate male *i*. Agrees to co-display at $A_{ij}^{Sub}$ = 1, is ambivalent at $A_{ij}^{Sub}$ = 0.5, and rejects at $A_{ij}^{Sub}$ = 0. | $A_{ij}^{Sub} = \begin{cases} 1 & G_{ij}^{Sub} > 0 \\ 0.5 & G_{ij}^{Sub} = 0 \\ 0 & G_{ij}^{Sub} < 0 \end{cases}$ |
| (14) | $A_{ij}^{Dom}$ | Decision by dominant male *j*. Agrees to co-display at $A_{ij}^{Sub}$ = 1, is ambivalent at $A_{ij}^{Sub}$ = 0.5, and rejects at $A_{ij}^{Sub}$ = 0. | $A_{ij}^{Dom} = \begin{cases} 1 & G_{ij}^{Dom} > 0 \\ 0.5 & G_{ij}^{Dom} = 0 \\ 0 & G_{ij}^{Dom} < 0 \end{cases}$ |

Preprint

**Appendix B**

*Ruff model Assumptions*

**The following specific assumptions are added to those described in Appendix A.**

1. Copulations are a valid measure of ruff reproductive fitness
2. Satellites do not gain copulations when displaying alone
3. A single competitive satellite is available for co-display at each lek
4. The presence of a satellite on a lek is equivalent to having another resident on the lek with the number of available copulations increasing accordingly
5. Copulations assigned to each resident are the copulations a resident can attract to his own court when a satellite is present on the lek but not co-displaying. Copulations are re-assigned when co-display occurs
6. Satellites will only choose one resident and co-display for time *T*, which is specified in the model (see Appendix B, *Time a satellite spends with preferred resident*)

Preprint

*Model parameterization*

*Resident mating skew*— This was taken from the graphical representation in figure 2b in Widemo and Owens (1995). However, since their reported regression formula did not match the plotted prediction lines (fig. B1), we digitalized the figure with R package “digitize” (Poisot, T. 2011) in r version (3.5.2 (2018-12-20) -- "Eggshell Igloo") and took 50 points along the prediction lines (Jpeg used for analysis is available at https://github.com/ruffresearcher5/co-display_in_lekking_ruffs/tree/main/supplementary_code/rcode_for_Widemo_and_Owens_fig_b2_analysis). We then estimated an exponential regression from those sample points. We adjusted the intercept of the model estimate so that the skew of a lek with a single resident would be equal to one. Similarly, for figure B2 we also extracted point predictions by hand (fig. B1).

```
library(digitize)

rskedata<-digitize(rske)#follow instructions, make 50 points along the trend line

rskem=nls(y~a-exp(b*x),data = rskedata,start=list(a=1,b=0.8))#negative exponential regression of digitized trend line

summary(rskem)#parameter estimates for regression, we adjusted the intercept estimate so that the skew at a lek size of one would be one

Formula: y ~ a - exp(b * x)


Parameters:

   Estimate Std. Error t value Pr(>|t|)

a 2.1059377 [adjusted to 2.06 in our model]  0.0039881   528.1   <2e-16 ***

b 0.0566548  0.0004289   132.1   <2e-16 ***

---

Signif. codes:  0 ‘***’ 0.001 ‘**’ 0.01 ‘*’ 0.05 ‘.’ 0.1 ‘ ’ 1


Residual standard error: 0.01488 on 57 degrees of freedom


Number of iterations to convergence: 11

Achieved convergence tolerance: 7.849e-06
```

Preprint

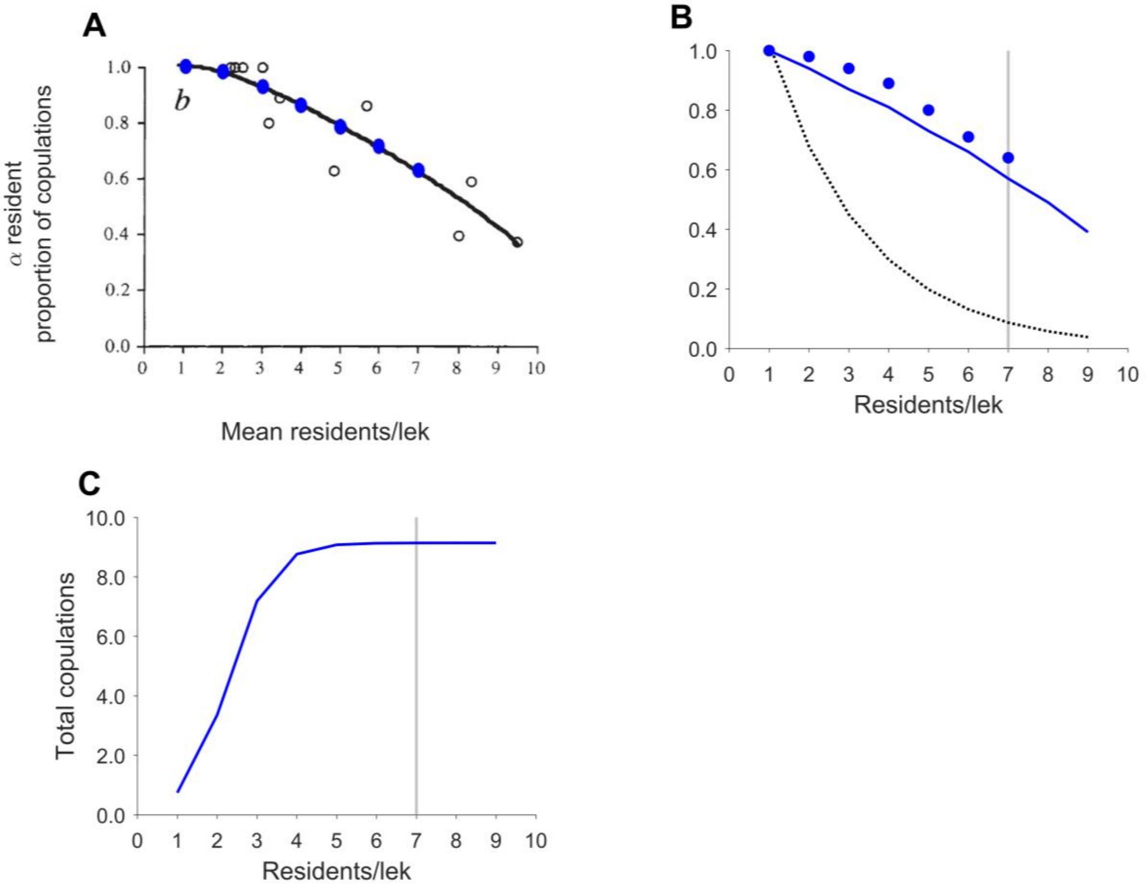


Figure B1. ***A*** Extracted by hand point predictions (blue closed circles) for each lek size from fig. 2b in Widemo and Owens (1995). Open circulars are original raw data from 15 leks. ***B*** Resident mating skew (proportion of copulations obtained by the α-resident) related to lek size. Blue line is adapted prediction line inferred from 50 digitized points, blue closed dots are extracted points from **A** and the black broken line is their regression ("'skew' = 1.5419 (0.6629$^{\text{'mean lek size'}}$"; fig. 2B, Widemo and Owens (1995)). *C* Total copulations available for a given lek size according to extracted values from 15 Swedish ruff leks (adjusted from Widemo and Owens 1995). Vertical grey lines in ***B*** and ***C*** indicate the maximum number of resident/lek in our model.

*Time a satellite spends with preferred resident and total copulations on a lek*—To estimate the mean time a satellite spends with his preferred resident during the peak lekking period, *T*, we took the observation time during the peak lekking season (10 days) from Widemo and Owens (1995) and combined this with the provided time that specified how long individual satellites codisplayed with their preferred resident (Table 4, van Rhijn 1991). We used the observation time frame from Widemo and Owens (1995) because the skew and copulation estimates we used came from this source. We extracted the following data from van Rhijn (1991) table 4: 1) percent time satellites spent with preferred resident (White-1 = 66%, white-2 = 40%, little-boy = 52%), and 2) the time in minutes satellites spent on the lek (White-1 = 139, white-2 = 53, little-boy = 61). We then multiplied the mean percent time a satellite spent with their preferred resident ($52.\overline{66}$ %) times the mean time a satellite spent on a lek $84.\overline{33}$ min/day. This was the mean time a satellite displayed with a preferred resident per day. We then multiplied the mean time a satellite displayed with a preferred resident per day with ten (the duration of the peak lekking season). Therefore, $T$, in our model was the mean time for co-display with the preferred resident during the observation period = 44.42 min/day • 10 days = 444.2 ~ 444 min.

*Fitness loss assessment*—Co-display on a lek leads to a fitness loss $H_r$ for each single-displaying resident $r$. The specific mechanism by which co-displays diverts copulations is unclear. To account for this lack of empirical data we examined four hypothetical fitness loss scenarios (table B1, number 3; fig. B2 and B3). Under, the “Null” effect scenario, co-displays leaves the copulation distribution of all residents on the lek the same ($H_r = 0$). In the other three scenarios copulations are re-distributed among resident courts ($H_r > 0$). Under the “Uniform Proportion” scenario, co-display draws the same proportion of copulations from each resident (table B1, number 4; fig. B2*B*). Under the “Skew” scenario, co-display draws a higher proportion of copulations away from high rather than low ranked residents. The exact proportion is based on the reproductive skew (table B1, number 5; fig. B2*C*). Under the “Reverse Skew” scenario, co-display draws the highest proportion of copulations from low rather than high ranked residents (table B1, number 6; fig. B2*D*).

We used all four models to assess which fitness loss scenario was most likely to occur in nature. Since the satellite allele is homozygous lethal (Küpper et al. 2016), we assumed that satellite copulations within co-displays should meet or exceed satellite copulation rates observed in the wild (~10%, Widemo 1998). Otherwise the satellite allele would go extinct. Hence, if the model predicted co-display to occur and the satellite reward $G_r^{Sat}$ was larger than $F^{Lek} \cdot 10\%$, we considered the scenario as biologically plausible.

Preprint

Figures B4 – B7, and D6 show the predictions of our model under the four hypothetical fitness loss scenarios. The “Null” scenario did not produce any co-display. In contrast, the “Uniform Proportion”, “Skew”, and “Reverse Skew” scenarios produced co-displays that are likely to occur in nature (fig. B7). However, under the “Reverse Skew” scenario co-display was only plausible on small lek sizes, which does not match observations in the wild (Höglund et al. 1993). Similarly, the “Uniform proportion” scenario did not suggest that co-display would not be viable on large leks as satellites failed to obtain enough copulations. Therefore, we present the “Skew” scenario in the main text. We found further support for the “Skew” effect in our sensitivity analysis, in Appendix D. This analysis shows that the “Skew” scenario allows for more variation in satellite competitive ability against their co-displaying resident. The “Skew” scenario predicted multiple realistic co-displays under each lek size for all levels of satellite competitive ability whereas the “Uniform Proportion” scenario only predicted one realistic co-display over all lek sizes when Satellites had strong competitive abilities (fig. D6).

Preprint

Table B1. Notation and definitions of different fitness loss scenarios. The “Skew” scenario was the scenario with the highest biological plausibility and is presented in the main text.

| Number | Elements | Description | Formula/value ranges |
| --- | --- | --- | --- |
| (1) | $h$ | Fitness loss scenario index. | $h \in [null, UP, Sk, RSk]$ |
| (2) | $H_r^h$ | Fitness loss, the proportion of copulations diverted from a resident’s court by a co-display[a]. | $H_r^h = \begin{cases} H_r^{null} \\ H_r^{UP} \\ H_r^{Sk} \\ H_r^{RSk} \end{cases}$ |
| (3) | $H_r^{null}$ | Fitness loss under the “Null” scenario. | $H_r^{null} = 0$ |
| (4) | $H_r^{UP}$ | Fitness loss under the “Uniform Proportion” scenario[b]. | $H_r^{UP} = \frac{B}{L}$ |
| (5) | $H_r^{Sk}$ | Fitness loss under the “Skew” scenario. | $H_r^{Sk} = B^2(1 - B)^{r-1}$ |
| (6) | $H_r^{RSk}$ | Fitness loss under the “Reverse Skew” scenario. | $H_r^{RSk} = B^2(1 - B)^{L-r}$ |

[a] $r$ is a subscript: describes a specific resident rank (α through η) and a variable: numeric equivalent of the $r$ subscript (e.g., when $r$ as a subscript = α, $r$ as a variable = 1).

[b] $B$ is the resident mating skew given lek size $L$.

Preprint

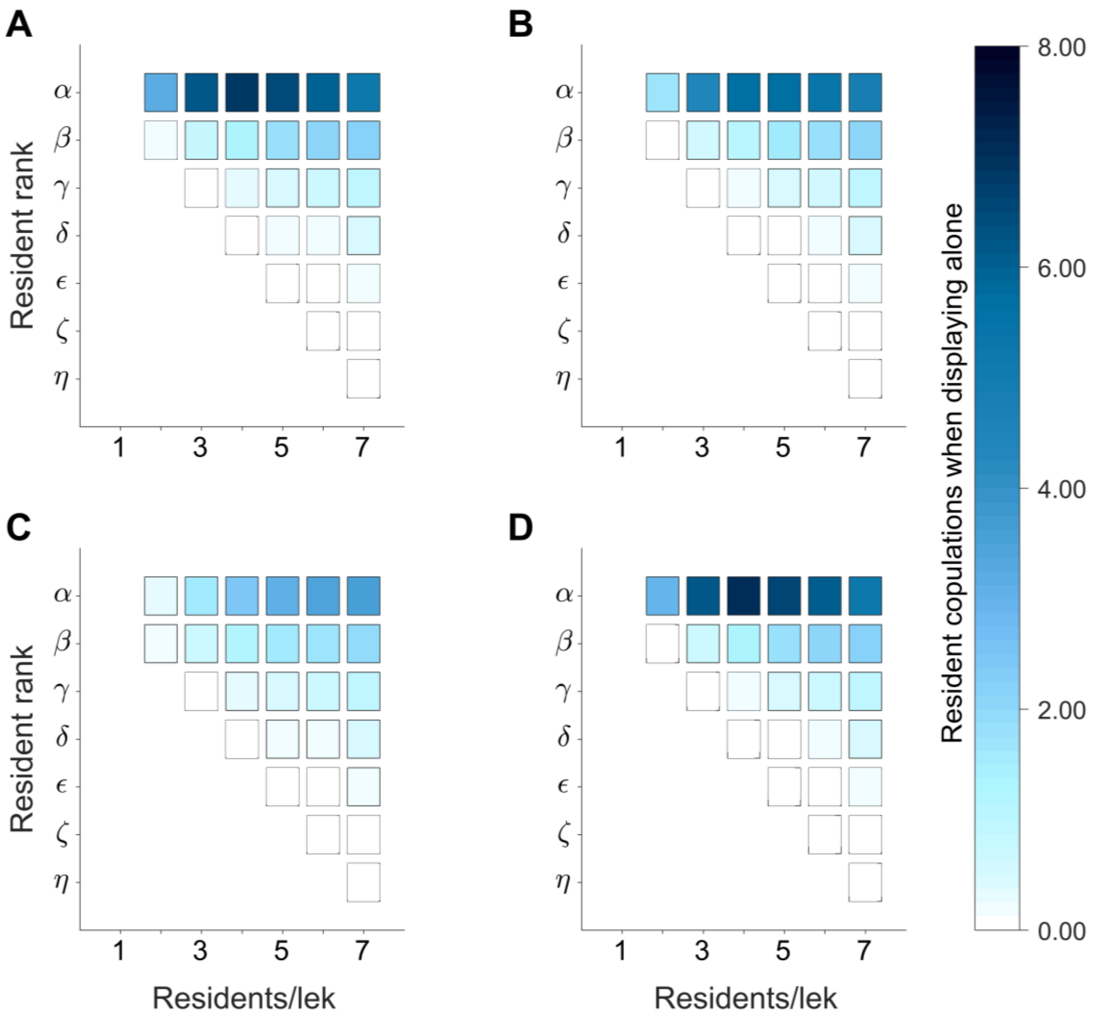


Figure B2. Resident copulations when a satellite is co-displaying with another resident on the lek $F_r^{Res}$. **A – D** represent different scenarios for the fitness loss $H_r$ for residents that display alone (see text). ***A*** Null scenario. ***B*** Uniform Proportion scenario. ***C*** Skew scenario. ***D*** Reverse Skew scenario. Note that the fitness loss is restricted to lek sizes $2 \leq L \leq 7$.

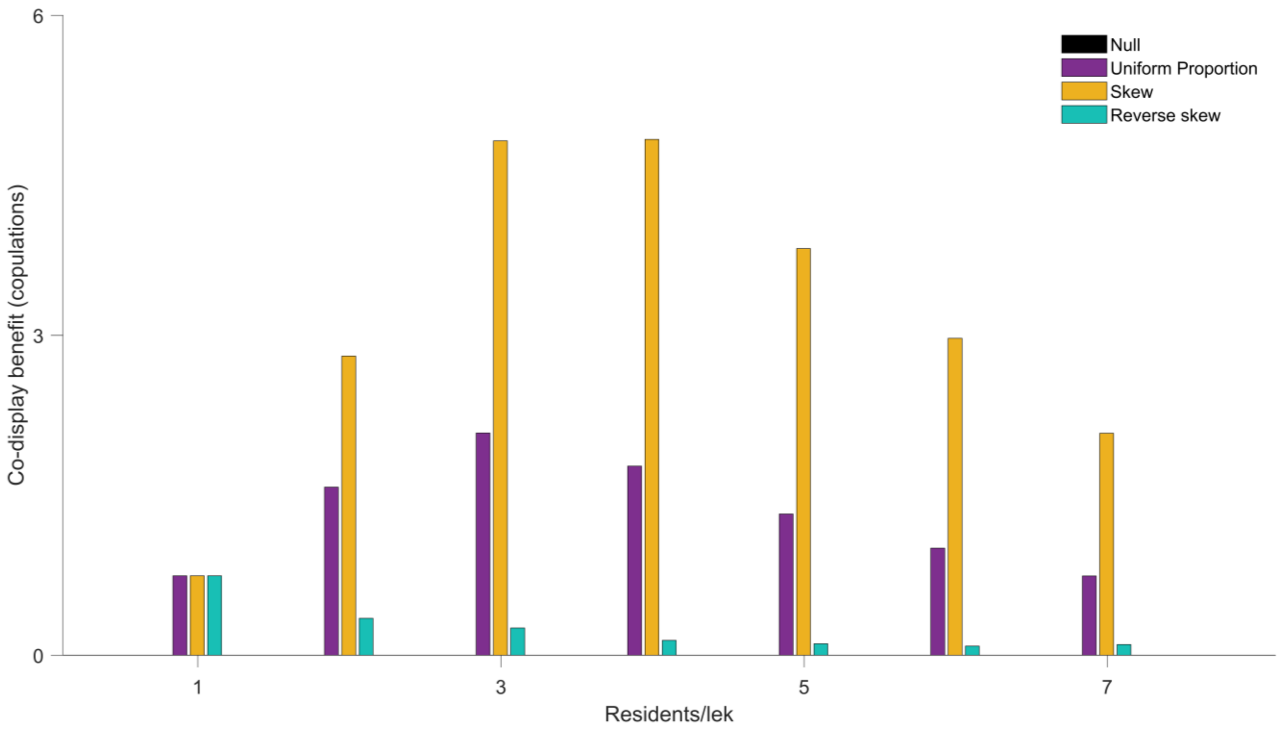


Figure B3. Co-display benefit $F^{Sat+Res}$ available to a co-displaying court depending on lek size and fitness loss scenarios, Null, Uniform Proportion, Skew, and Reverse Skew. Note that in the Null Scenario, in black, $F^{Sat+Res} = 0$, and is therefore not visible on the figure.

Prepr

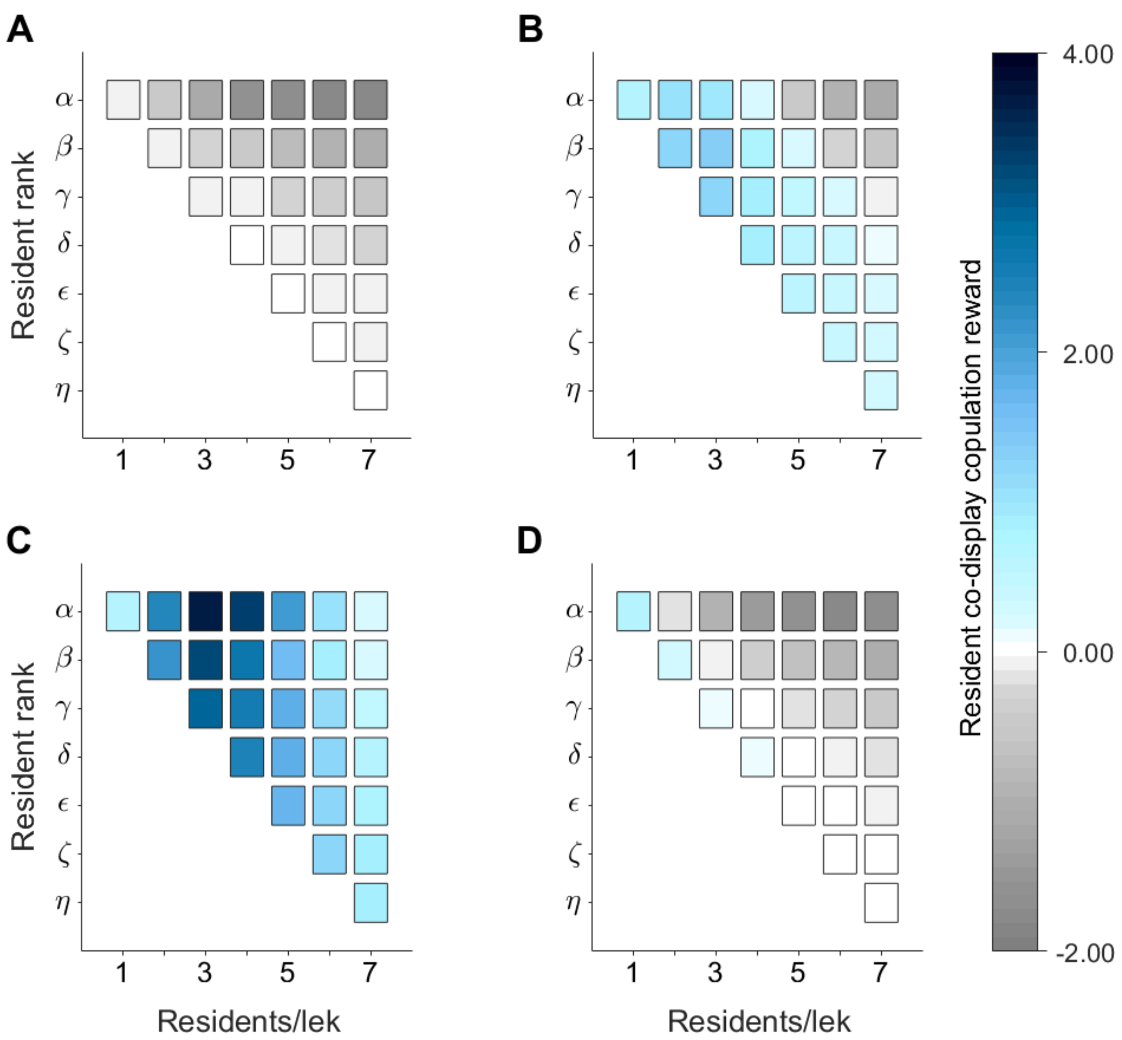


Figure B4. Resident copulation reward $G_r^{Res}$ according to resident's rank, lek size, and fitness loss scenario. $G_r^{Res}$ determines whether a resident will co-display or not. Residents reject satellites when their copulation gains are less than zero (grey) and agree to co-display with them when gains are greater than zero (blue). Note that very small values (both negative and positive) appear white as they approach zero. ***A*** Null scenario, ***B*** Uniform Proportion scenario, ***C*** Skew scenario, and ***D*** Reverse Skew scenario.

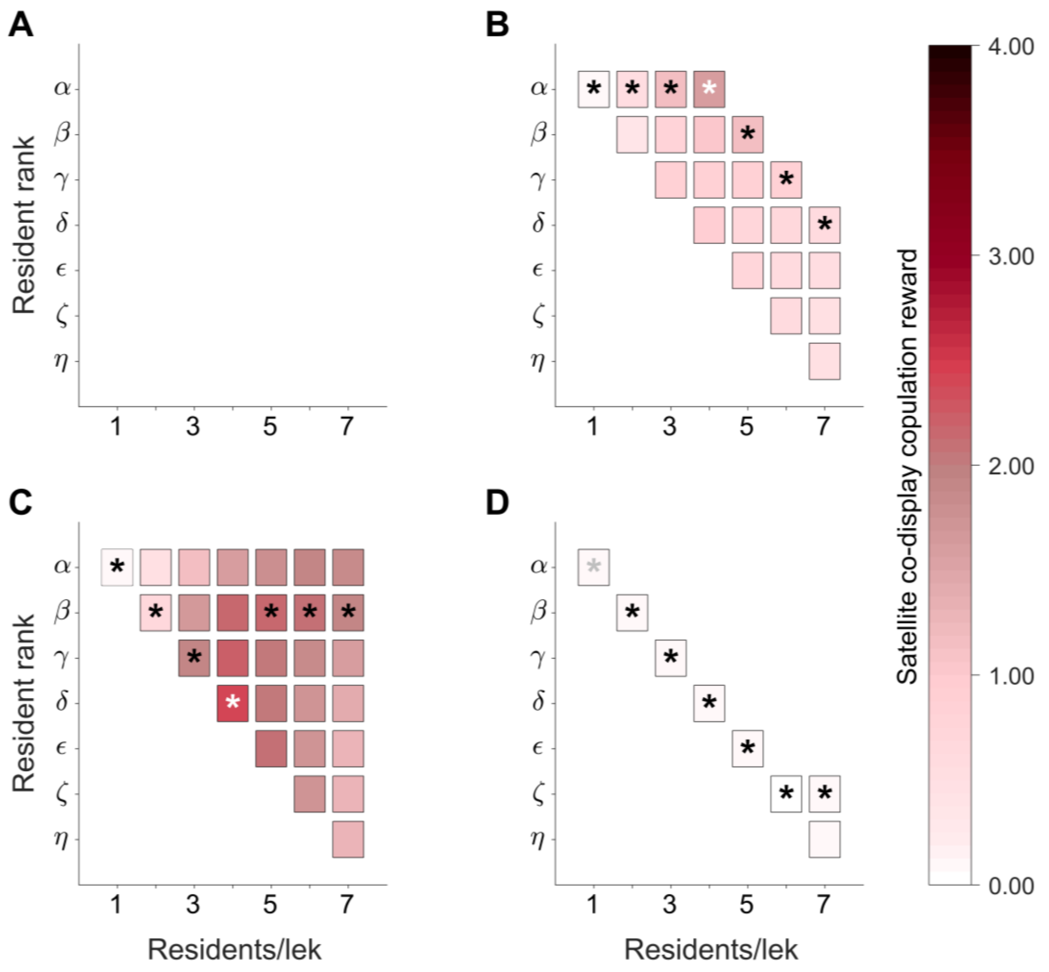


Figure B5. Satellite copulation reward $G_r^{Sat}$ according to the rank of his resident partner, lek size and fitness loss scenario. Missing squares refer to resident/satellite combinations where co-display is predicted not to occur. Asterisks indicate preferred choice at a given lek size. White/gray asterisks, indicate where satellites achieve the highest copulation reward over all lek size within each scenario. ***A*** Null scenario, ***B*** Uniform Proportion scenario, ***C*** Skew scenario, and ***D*** Reverse Skew scenario.

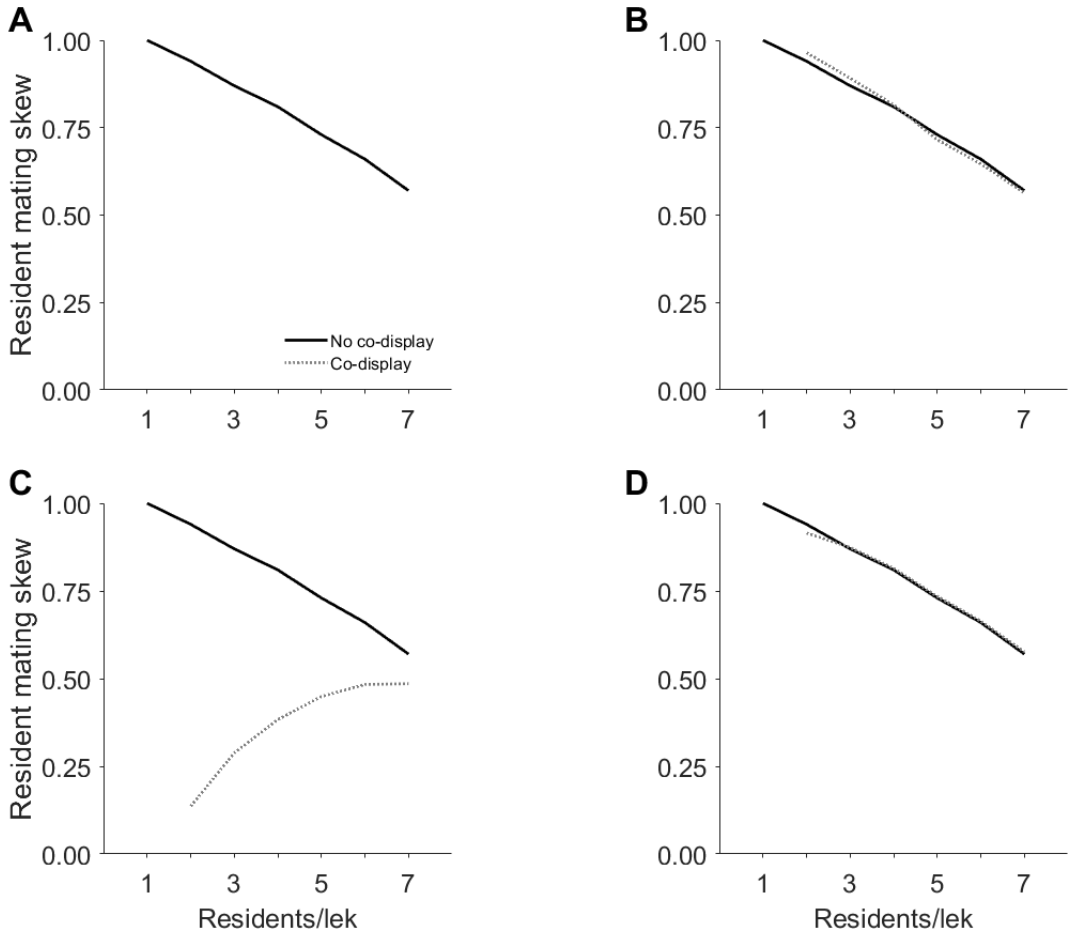


Figure B6. Mating skew among residents with or without co-display according to lek size and fitness loss scenario. Shown is the co-display with the preferred partner of the satellite at a given lek size taken from Fig. B5. ***A*** Null scenario, co-display never occurs and the mating skew remains the same. ***B*** Uniform Proportion scenario, co-display increases skew at lower lek sizes but decreases it at larger lek sizes. ***C*** Skew scenario, co-display always reduces the mating skew. ***D*** Reverse Skew scenario, at all lek sizes, except $L$ = 2, the mating skew remains the same with and without co-display.

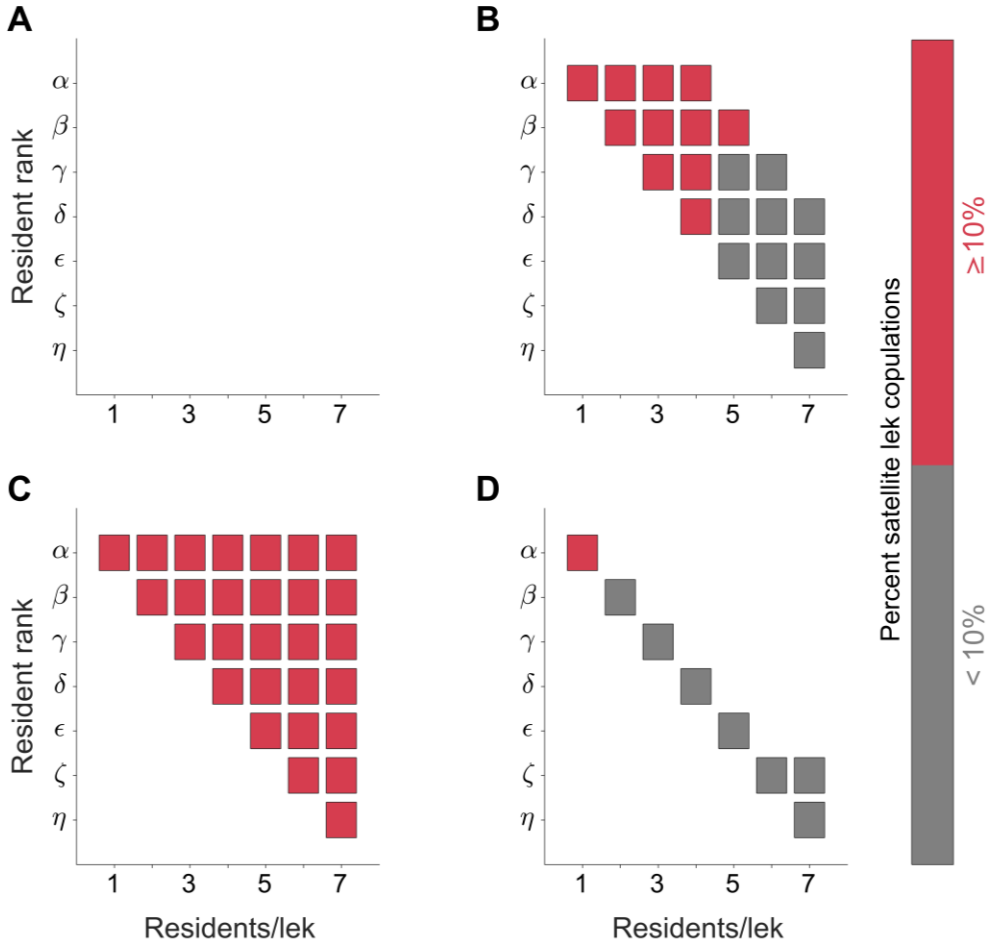


Figure B7. Proportion of copulations obtained by a satellite co-displaying with a given resident, by resident's rank, lek size, and fitness loss scenario. Combinations in red are biologically plausible, whereas those in grey are not. Missing squares refer to resident/satellite combinations where co-display is predicted not to occur. ***A*** Null scenario, ***B*** Uniform Proportion scenario, ***C*** Skew scenario, and ***D*** Reverse Skew scenario.

*Resource holding potential of residents and disruption risk relationships*—

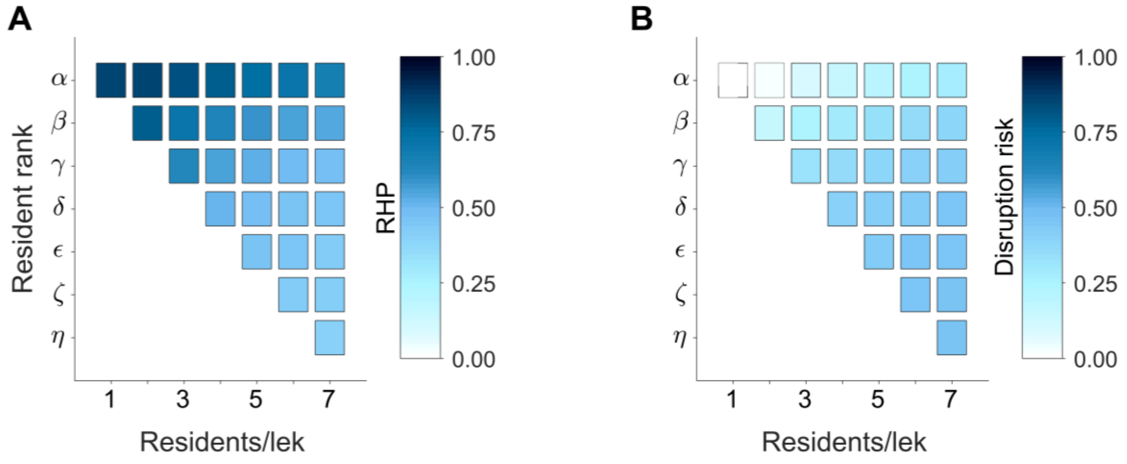


Figure B8. Model assumption inputs for the resident/satellite co-display model according to lek size. ***A*** Resource holding potential (RHP) of residents for each hierarchical rank. ***B*** Disruption risk of residents, specifying the rank-dependent probability that a copulation by a resident will be interrupted by another resident.

Preprint

### Appendix C

*Limits of Resident co-display choice*

The limits of cooperation by a resident can be expressed in terms of the resource holding potential (RHP) $M_r^{Res}$ of a given resident and the fitness loss $H_r$. A resident compares his copulations when displaying alone to those when he is co-displaying to decide to join a satellite in co-display. The difference between a resident *r*'s copulations through co-display and single display is then expressed by:

$$M_r^{Res}(F_r^{Res} + F^{Sat+Res}) - (1 - H_r)K_r$$

We can then substitute in the definitions of $F_r^{Res}$ and $F^{Sat+Res}$:

$$F_r^{Res} = (1 - H_r)K_r$$

$$F^{Sat+Res} = \sum_{r=\alpha}^{L} H_r K_r$$

So, the difference becomes:

$$M_r^{Res}\left((1 - H_r)K_r + \sum_{r'=\alpha}^{L} H_{r'}K_{r'}\right) - (1 - H_r)K_r$$

Here $r'$ is an index for any resident while *r* is an index for the focal resident. We can split the sum:

$$M_r^{Res}\left((1 - H_r)K_r + H_rK_r + \sum_{r' \neq r} H_{r'}K_{r'}\right) - (1 - H_r)K_r$$

$$M_r^{Res}\left(K_r + \sum_{r' \neq r} H_{r'}K_{r'}\right) - (1 - H_r)K_r$$

$$M_r^{Res}K_r - (1 - H_r)K_r + M_r^{Res}\sum_{r' \neq r} H_{r'}K_{r'}$$

$$\left(M_r^{Res} - (1 - H_r)\right)K_r + M_r^{Res}\sum_{r' \neq r} H_{r'}K_{r'}$$

Preprint

The resident will always cooperate (agrees to co-display/ambivalence) when this difference is equal to or greater than zero:

$$\left(M_r^{Res} - (1 - H_r)\right)K_r \; + \; M_r^{Res} \sum_{r' \neq r} H_{r'} K_{r'} \geq 0 \xrightarrow{yields} \text{Resident cooperates}$$

since $K_r$ and the sum are positive, when:

$$M_r^{Res} - (1 - \; H_r) \; \geq 0$$

$$M_r^{Res} \geq 1 - H_r$$

In words, the residents accepts the satellite whenever his RHP exceeds the fraction by which his copulations are diminished when the satellite co-displays with another resident.

Preprint

**Appendix D**

*Sensitivity to satellite competitive ability and disruption risk*

Table D1. Elements of the sensitivity analysis of the ruff resident/satellite cooperation model described in the main text.

| Number | Elements | Description | Formula/value ranges |
|---|---|---|---|
| (1) | $M^{Sat}$ | Satellite competitive ability (see Fig. D1). Satellites had three competitive abilities: weak, medium, and strong. Weak satellites only gain copulations when a co-displaying resident competes with other residents. Satellites of medium ability could obtain at least 15% lek when paired with the α-resident on a small lek whereas strong satellites could obtain 30% under the same conditions. | $M^{Sat} = \begin{cases} 0.00 & \text{weak} \\ 0.15 & \text{medium} \\ 0.30 & \text{strong} \end{cases}$ |
| (2) | $D_r$ | Disruption risk (see Fig. D2). There were three competition levels that altered disruptive risk: low competition, where residents interrupt each other infrequently; medium competition, residents interrupt each other with a medium frequency; and high competition, residents are interrupted almost 50% percent of the time. | $D_r = \begin{cases} \frac{1}{1+e^{\left(\frac{1}{Lr}\right)\cdot 14}} & \text{low} \\ \frac{1}{1+e^{\left(\frac{1}{Lr}\right)\cdot 7}} & \text{medium} \\ \frac{1}{1+e^{\left(\frac{14}{Lr}\right)\cdot 7}} & \text{high} \end{cases}$ |

Preprint

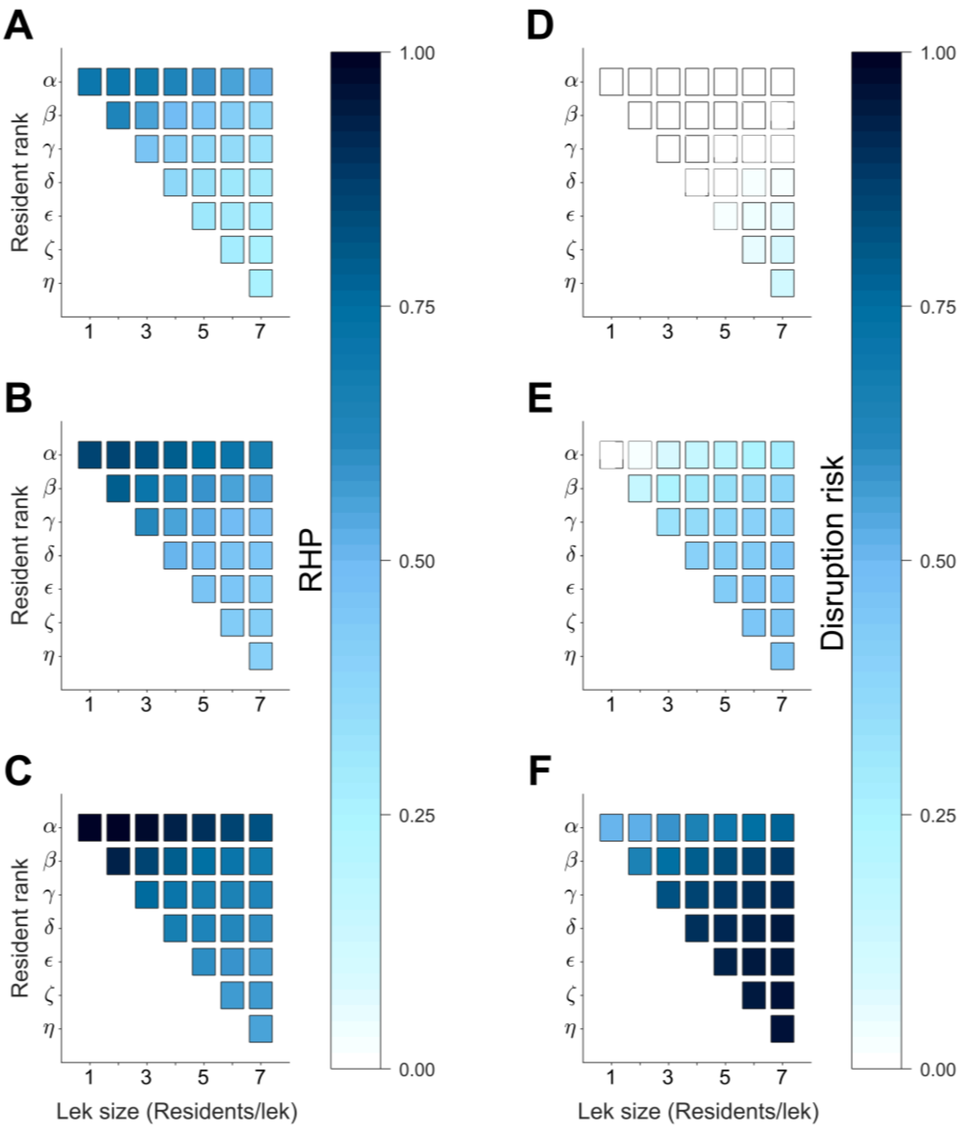


Figure D1. Variation of resource holding potential and disruption risk of residents according to lek size and hierarchical rank for sensitivity analysis. ***A* – *C*** Monopolizations coefficients when satellites differ in their competitive ability: ***A*** strong satellite, ***B*** medium satellite, ***C*** weak satellite. ***D* – *F*** Disruptive risk when competition between residents differs: ***D*** low competition, ***E*** medium competition, ***F*** high competition.

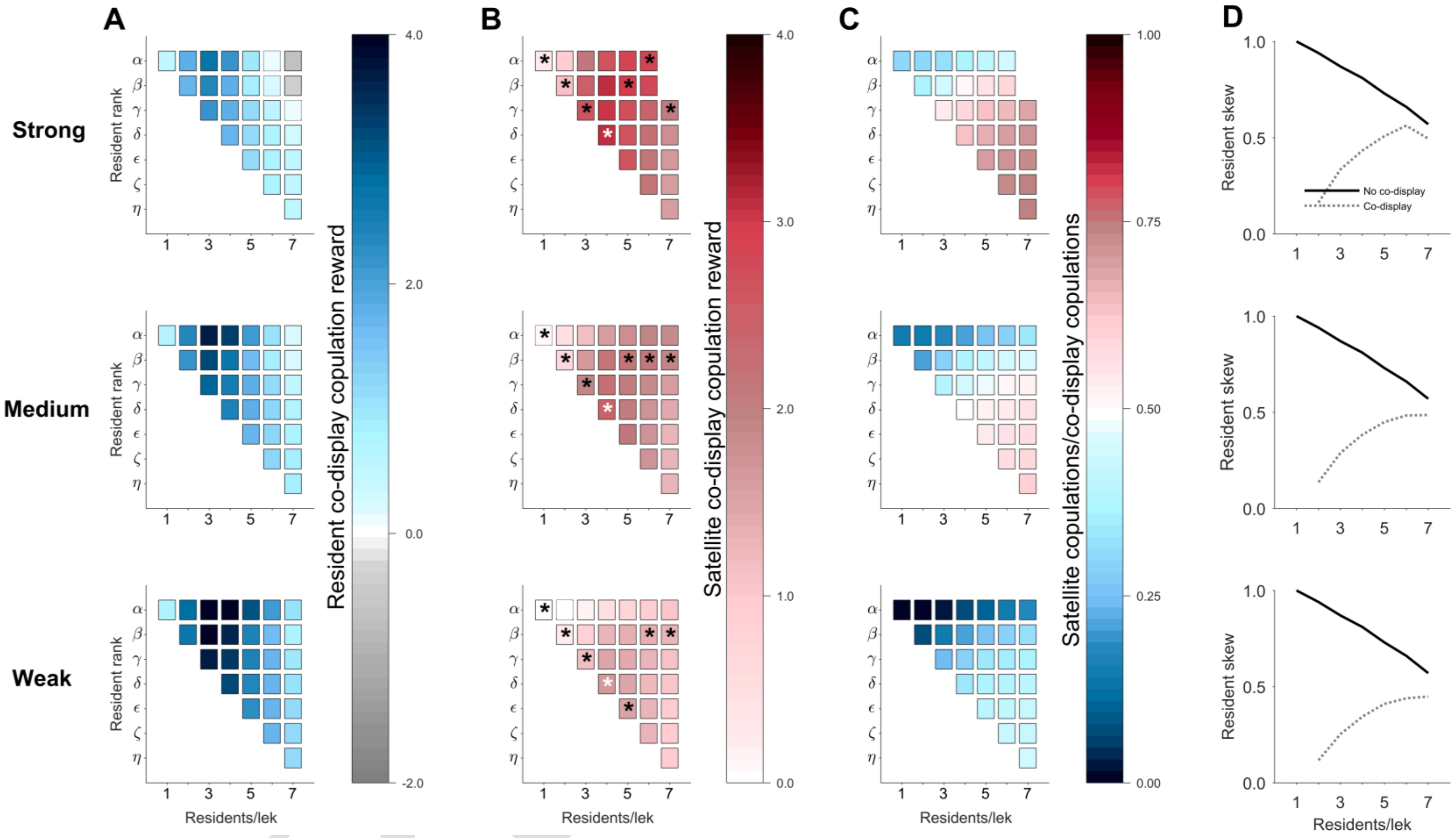


Figure D2. Sensitivity of resident/satellite co-display model to variation in satellite competitive ability (“Strong”, “Medium”, “Weak”). ***A*** Resident copulation reward from co-display. ***B*** Satellite copulation reward from co-display. Asterisk refers to chosen resident partner at a given lek size. White asterisk represents overall preferred resident. ***C*** Satellite’s proportion of copulations from co-display. ***D*** Resident mating skew. Grey or missing squares mean that co-display does not occur.

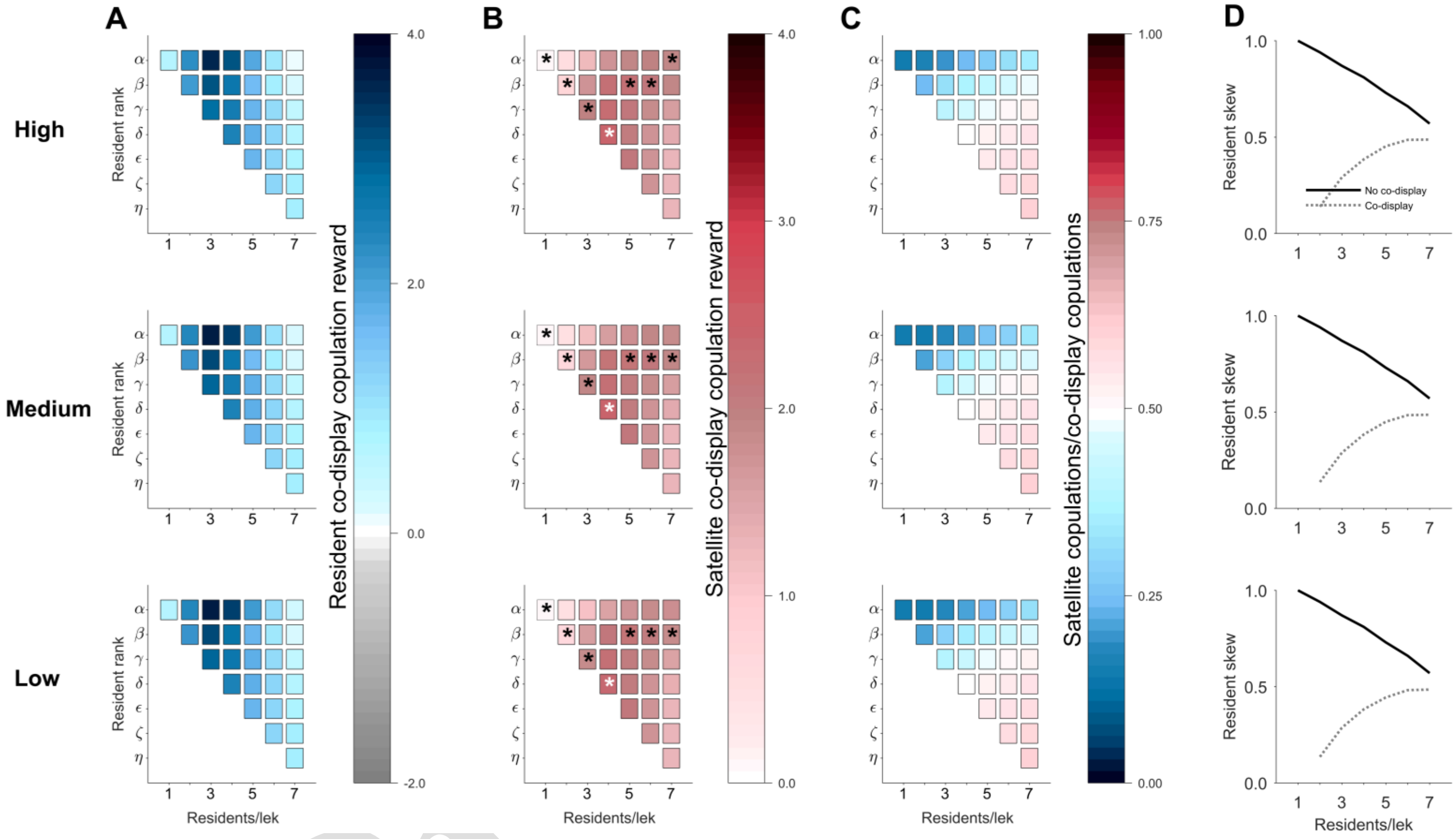


Figure D3. Sensitivity of resident/satellite co-display model to variation in disruptive risk ("High", "Medium", "Low"). ***A*** Resident copulation reward from co-display. ***B*** Satellite copulation reward from co-display. Asterisk refers to chosen resident partner at a given lek size. White asterisk represents overall preferred resident. ***C*** Satellite's proportion of copulations from co-display. ***D*** Resident mating skew.

PJ

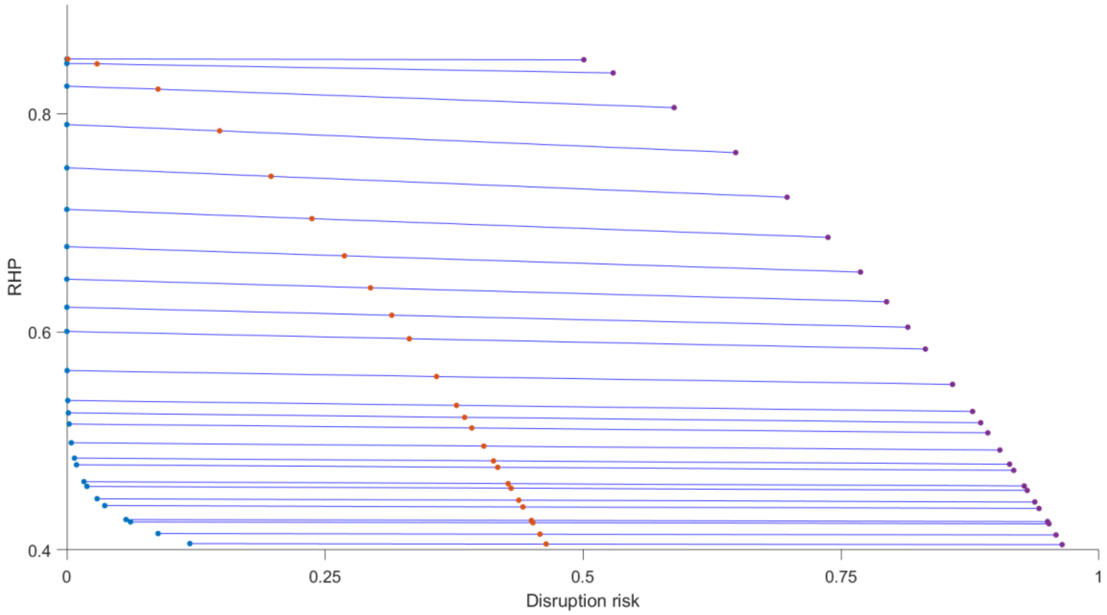


Fig. D4. Relationship between the resource holding potential of residents (RHP) and disruption risk for all leks sizes ($L \in [1, 7]$) and hierarchical ranks ($r \in [\alpha,\eta]$) combined. Blue circles are a resident's RHP predicted by their disruption risk when there is little competition on the lek. Orange circles indicate medium competition levels on the leks. Purple circles represent high competition on leks. Blue lines indicate a minor decrease in RHP due to increases in disruption risk.

Preprint

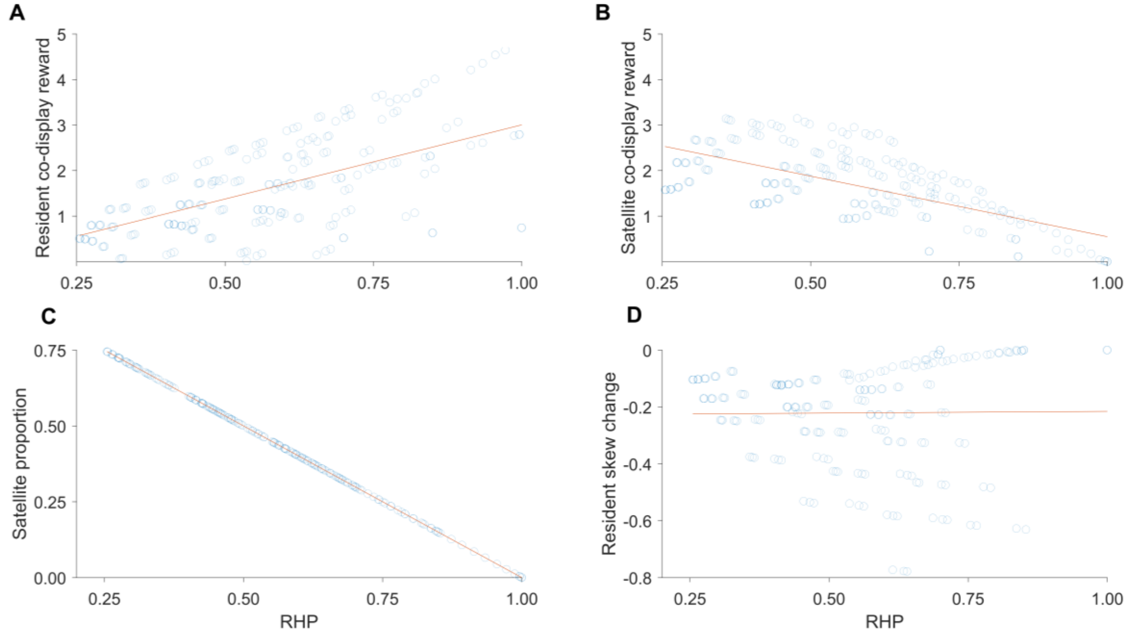


Fig. D5. Influence of the resource holding potential (RHP) of the resident on fitness parameters in satellite/resident co-display model. Blue curricles are responses to perturbed RHP and red lines are estimated linear regression lines. ***A*** Resident co-display reward is positively related to RHP (regression: $G_r^{Res} = -0.25735 + 3.2607M_r^{Res}$). ***B*** Satellite co-display reward is negatively related to RHP of his resident partner (regression: $G_r^{Sat} = 3.2043 - 2.6566M_r^{Res}$). (Note that changes in $G_r^{Res}$ and $G_r^{Sat}$ do not have the same absolute slope in relation to $M_r^{Res}$ instead the payoffs for the resident, $C_r^{Res}$, and the satellite, $C_r^{Sat}$, do. This is because $G_r^{Res} = C_r^{Res} - F_r^{Res}$ and $G_r^{Sat} = C_r^{Sat} - 0$.) ***C*** Satellite copulation proportion in dyad is negatively related to RHP of his resident partner (regression: $Sat.prop. = 1 - 1M_r^{Res}$; indicating that the payoffs of satellites and residents are proportional). ***D*** Resident mating skew is not related to RHP (regression: $Delta\ Res.skew. = -0.2274 + 0.0119M_r^{Res}$).

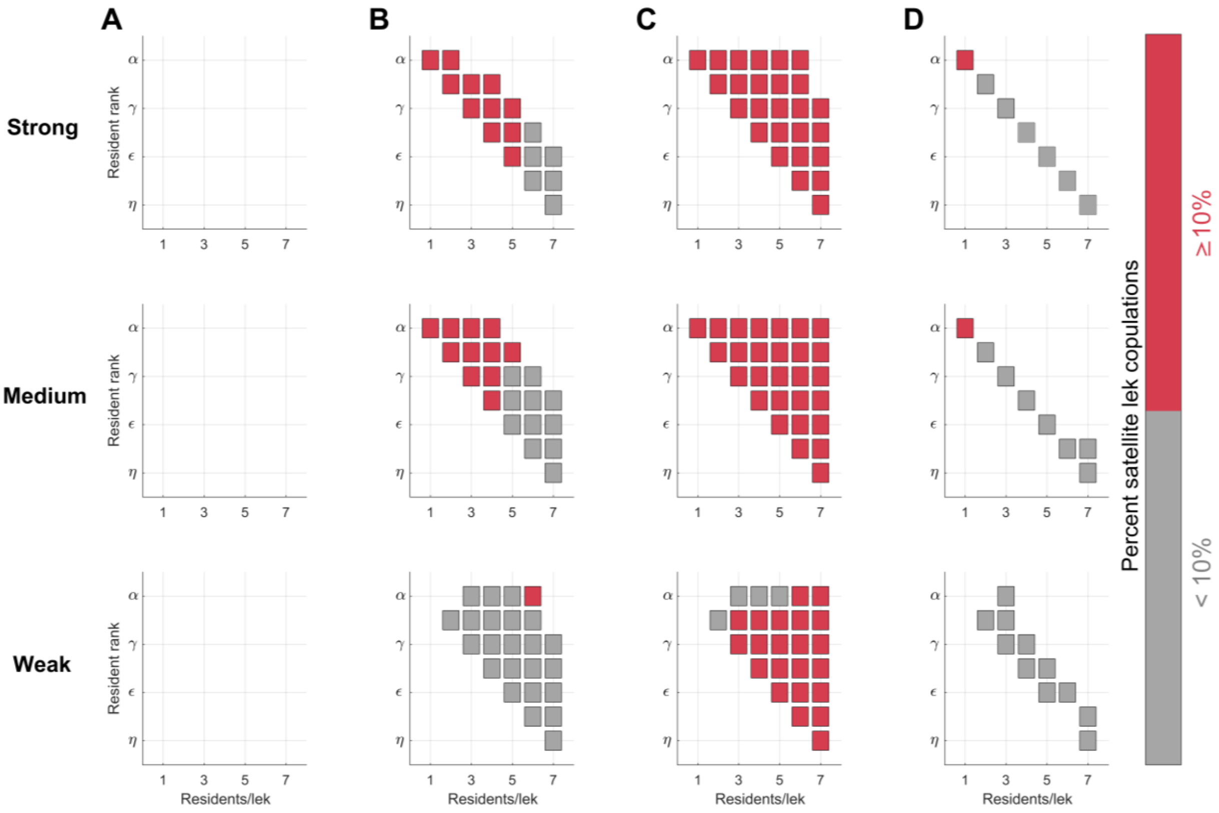


Figure D6. Sensitivity of the proportion of satellite copulations on a lek to variation in satellite competitive ability (“Strong”, “Medium”, “Weak”) under four different fitness scenarios. Satellites require at least 10% of copulations to avoid going extinct. ***A*** Null scenario, ***B*** Uniform Proportion scenario, ***C*** Skew scenario, ***D*** Reverse Skew scenario.

Pre

Preprint
